# Structural plasticity of 2A proteins in the Parechovirus family

**DOI:** 10.1101/2024.04.06.588386

**Authors:** Ling Zhu, Marion Pichon, Zuzanna Pietras, Xiangxi Wang, Jingshan Ren, Elizabeth E. Fry, David. I. Stuart, Anastassis Perrakis, Eleonore von Castelmur

## Abstract

Parechoviruses, including *Parechovirus A* that infects humans as well as *Parechovirus B* (formerly Ljungan virus) and *Parechovirus C* (formerly Sebokele virus) that infect rodents, belong to a group of picornaviruses whose 2A proteins, instead of being proteases, contain a conserved H-box and NC-motif and are homologous to a small cellular lipid-modifying enzyme (PLAAT3) that acts as a host factor, enabling the picornavirus life cycle. Despite the common evolutionary origin, 2A^H/NC^ proteins and PLAAT3 have no conserved function, as the active site of the viral proteins cannot support catalysis. Here, we set out to find if all *Parechovirus* species share the structural rearrangement that destroys the active site configuration of the cellular enzyme. This has revealed a remarkable structural plasticity of these 2A^H/NC^ proteins that arises not only from sequence differences between species, but also from differences in the length of the recombinantly expressed proteins, resulting in large structural rearrangements. These include rerouting of a large internal loop and repositioning of the C-terminal helix with respect to the central β-sheet, and these in turn influence the oligomeric state of the protein. We discuss how this structural plasticity could correlate with the function of these proteins in the viral life cycle and how this could recapitulate the possible evolution of this protein from host factor to viral 2A^H/NC^ protein, with new independent functions in RNA replication.

## Introduction

Human Parechoviruses (HPeVs) are members of the large and medically important family of picornaviruses. HPeV1 and HPeV2 were originally isolated over sixty years ago from children presenting with diarrhea and characterized as echoviruses 22 and 23, based on their morphology and clinical symptoms. Genetic and molecular characterization has since revealed many unique characteristics regarding their genome organization, structure, and replication, setting them apart from other viruses in this group. Therefore, they have been re-classified as the prototypic members of the species *Parechovirus A*, and assigned to a separate genus, *Parechovirus* (1, 2). Ljungan virus (LV), first isolated from bank voles (3), has since been classified as the type-member of a second species, *Parechovirus B*. It shares many of the unusual features seen in *Parechovirus A*, and was the first picornavirus that was shown to contain two distinct 2A proteins (denoted as 2A_1_ and 2A_2_) (4). The genomic characterization of Sebokele virus (SEBV1) shows that it is most closely related to Ljungan virus and also contains two distinct 2A proteins, but it is classified as a third species, *Parechovirus C* (5).

Recently, the crystal structures of HPeV1 (6) and cryo-EM structures of HPeV3 (7, 8) and of LV1 (9) have been reported. A structural comparison with other picornaviruses shows that they are most closely related to Hepatitis A virus (HAV; (10)) and places these viruses close to the insect picorna-like viruses, confirming previous phylogenetic analysis locating the parechoviruses basal to the picornavirus tree (5, 11, 12). The structural features characteristic of parechoviruses include i) the lack of a hydrophobic canyon, ii) the absence of a pocket factor, iii) the domain swapping of the N-terminus of VP0 across the icosahedral two-fold axis, iv) a C-terminal extension to VP1, which might be the short 2A_1_ protein(13). Furthermore, the structure of LV1 reveals that the basic extension at the N-terminus of VP3, which is clustered around the five-fold axis, interacts with the viral RNA, leading to the ordering of a substantial fraction of the genome. The HPeV1 structure also revealed density for ordered RNA around the five-fold axis, allowing the modelling of a hexanucleotide. The interaction of the genome with the capsid proteins hints at a possible role of the genome in the virion assembly – the highly charged proteins might not be able to assemble into a stable capsid in absence of RNA due to electrostatic repulsion. Furthermore, the lack of VP4, which (in other picornaviruses) is implicated in the pore formation crucial for viral uncoating and genome translocation, suggests that parechoviruses may also uncoat in a different manner. ((14, 15))

Parechoviruses, unlike most picornaviruses, encode 2A proteins that are not proteases involved in polyprotein processing or host protein translation shutdown. Rather, these 2A proteins, which were also shown to be involved in RNA replication ((16, 17)), are homologous to a cellular protein, PLAAT3 (18), sharing conserved sequence motifs, including the H-box and NC-motif, (which encompass the catalytic residues of PLAAT3) and are therefore called 2A^H/NC^ proteins. In Parechovirus B and C, 2A^H/NC^ is preceded by a short 2A_1_ protein of the 2A^npgp^ type, which leads to the separation of P1 from the rest of the polyprotein through a process called ribosome skipping (19). As we recently described, PLAAT3 is an essential host factor for picornaviruses (20), suggesting that these viruses carrying a H/NC-type 2A protein might have acquired and evolved this cellular protein to become independent of the host factor. However, structural characterization elucidated that a topological rearrangement in the H/NC-type 2A protein leads to an active site conformation incompatible with catalysis.

Here, we set out to find if different parechovirus species share the structural rearrangement that destroys the active site configuration observed in the cellular enzyme, by elucidating the crystal structures of several representative 2A^H/NC^ of the different parechovirus family members. We show that all of them retain the inactive structural configuration of the active site, and that this rearrangement appears to go hand in hand with higher order oligomeric assembly. Furthermore, our studies reveal that the C-terminal region appears to play an important role in regulating the structural plasticity and the oligomerization state of this protein. Further experiments are required to answer the still open questions how this structural plasticity correlates with function and whether it is linked to other unique features of the virus and its role(s) in the viral life cycle.

## 2. Materials and Methods

### 2.1 Cloning, Protein expression and purification

Codon optimised synthetic gene Blocks (GeneArt and Integrated DNA Technologies) were used for the cloning of HPeV1-2A protein (Uniprot ID Q66578: residues 774-902) and LV4-2A_2_ (strain 64-7855; Uniprot ID C0J6D4: residues 823-957) into the pETNKI 6xHis-3C-ORF bacterial expression vectors with either a Kanamycin or Ampicillin resistance cassette of the NKI Ligation Independent Cloning suite (21). The resulting constructs were fused N-terminally to the residues MAHHHHHHSAALEVLFQ-//-GPG, containing a HRV 3C protease cleavage site. LV4-2A_2_^1-122^ was generated by inserting a stop codon using the Quikchange method, following the manufacturer’s protocol. LV1-2A_2_ (isolate 87-012, NCBI Reference Sequence: NP_705878.1), HPeV3-2A (GenBank: BAC23086.1, residues 772-920) and SEBV1-2A_1_-2A_2_ (NCBI Reference Sequence: YP_008119838.1) were synthesized by the Genscript Company. Constructs for LV1-2A_2_ and HPeV3-2A expression in *E. coli* were cloned in pET28a vector between the Nco1 and Xho1 restriction sites. The resulting constructs contain the additional residues MS at the N-terminus, introduced during cloning prior to the protein sequence as well as a C-terminal, non-cleavable hexahistidine-tag. The construct for SEBV1-2A_2_ expression in *E. coli* was cloned in the pGEX-6P-1 vector between BamH1 and Xho1 restriction sites with a N-terminal GST fusion tag. During restriction digestion cloning residues GPLGS and LTRAAAS were introduced at the N- and C-terminus of the construct respectively. All clones were verified by DNA sequencing.

All plasmids coding for the different 2A proteins were transformed into *E.coli* BL21 (DE3) cells. The bacterial cultures were grown until OD ≈ 0.6 and protein expression was induced using 0.25-0.5 mM Isopropyl β-D-1-thiogalactopyranoside (IPTG) and cultures were grown overnight at 16-18°C (120 rpm). After harvesting, the cells were resuspended into cold lysis buffer containing 20 mM HEPES pH 7.4, 150-300 mM NaCl and 2mM TCEP prior to sonication. Lysates were clarified by high-speed centrifugation prior to the affinity chromatography step. SEBV1-2A_2_ was purified by binding to glutathione Sepharose 4B resin (Cytiva) and eluted with lysis buffer supplemented with 10mM reduced glutathione (GSH). The other 2A proteins were separated from the lysate using immobilized metal affinity chromatography with either TALON beads (Clontech) or chelating Sepharose beads charged with NiCl_2_ (Cytiva), employing a gradient of imidazole (20-250 mM) supplemented in the lysis buffer for the elution step. PreScission Protease (Cytiva) was added for the cleavage of the N-terminal GST tag (4°C, 6h) of SEBV1-2A. For the His_6_-tag tagged proteins, tag removal was by cleavage with 3C-HRV protease in conjunction with a dialysis step against 20 mM HEPES pH 7.4, 150 mM NaCl, 2 mM TCEP. After subtractive affinity purification, a size exclusion chromatography step was carried out using HiLoad Superdex 16/60 75 pg or 200 pg (SEBV1-2A_2_)(Cytiva), equilibrated with 20 mM HEPES pH 7.4, 150 mM NaCl and 2mM TCEP. Fractions containing the pure protein were pooled together, concentrated, flash-frozen in liquid nitrogen and stored at −80°C until required. The purifications of LV1-2A, HPeV3-2A and SEBV1-2A_2_ for crystallization were carried out without the addition of TCEP into the different buffers.

For selenomethionine incorporation, the SelenoMet^TM^ Medium (Molecular Dimensions Limited) was used according to the manufacturer’s instructions for protein expression in the methionine auxotrophic *E. coli* strain 834 (DE3). Protein purification followed the same protocol as for the native proteins.

### 2.2 Crystallization

Crystallization was by vapor diffusion in a sitting drop plate setup for all proteins. Crystals of HPeV3-2A (70 mg/ml) were grown at 20°C in Greiner CrystalQuick X plates in 1.7 M Ammonium sulphate, 0.05 M Sodium Cacodylate pH 6.5 by mixing 100 nl protein with 100nl reservoir solution. Crystals were transferred into a solution containing mother liquor supplemented with 20% (v/v) glycerol and vitrified in liquid nitrogen for diffraction studies. Crystals of LV1-2A_2_ (75 mg/ml) were grown at 20°C in Greiner CrystalQuick X plates from 20% PEG3350, 0.2 M sodium formate by mixing 100 nl protein with 100 nl reservoir solution. Crystals were transferred into a solution containing mother liquor supplemented with 20% (v/v) glycerol and vitrified in liquid nitrogen for diffraction studies. Crystals of LV4-2A_2_^1-122^ (∼30.3mg/ml) were grown in MRC 2-well sitting drop plates. Crystal form 1 (I222) grew at 4°C in in 0.1M Na acetate pH 5.25, 10% PEG 6000 by mixing 100 nl protein with 100 nl reservoir solution. Crystal form 2 (P2_1_2_1_2_1_) grew at room temperature in 0.1M Tris pH8.0, 8% PEG 6000, 0.15M NaCl by mixing 200 nl protein with 100 nl reservoir solution. Crystals were transferred into a solution containing mother liquor supplemented with 25% (v/v) PEG 400 and vitrified in liquid nitrogen for diffraction studies. Crystals of SEBV1-2A_2_ (40 mg/ml) grew at 20°C in CrystalQuick X plates from 30% PEG4000, 0.005 M magnesium acetate, 0.050 M sodium cacodylate pH 6.5. Crystals were transferred into a solution containing mother liquor supplemented with 20% (v/v) glycerol and vitrified in liquid nitrogen for diffraction studies. Crystals of HPeV1-2A^1-129^ were grown in MRC 2-well sitting drop plates. Crystals of native HPeV1-2A^1-129^ (∼24.3 mg/ml) were grown in condition A4 of the ComPAS screen (NeXtal Biotech) containing 15% PEG 8000, 0.5M lithium sulphate by mixing 200 nl protein with 100 nl reservoir solution. Crystals of selenomethionine substituted HPeV1-2A^1-129^ (23mg/ml) were obtained in condition B11 of the Procomplex screen (NeXtal Biotech) containing 0.1 M Hepes-NaOH pH7.0, 15% PEG 4000 by mixing 100 nl protein with 100 nl reservoir solution. Crystals were transferred into a solution containing mother liquor supplemented with 25% (v/v) ethylene glycol and vitrified in liquid nitrogen for diffraction studies.

### 2.3 X-ray diffraction data collection, data processing, structure solution and refinement

Diffraction data for HPeV3-2A, LV1-2A_2_ and SEBV1-2A_2_ were collected at beamline I03 at Diamond Light Source (DLS) and processed using Xia2 (22). Diffraction data for native HPeV1-2A^1-129^ were collected at beamline ID 23-1 at the European Synchrotron Radiation Facility (ESRF), while diffraction data for selenomethionine labelled HPeV1-2A^1-129^ and LV4-2A_2_ ^1-122^ were collected at beamline PX3 at the Swiss Light Source (SLS), integrated with the XDS package(23) and scaled and merged using AIMLESS (24).

Phasing for HPeV3-2A, LV1-2A_2_ and LV4-2A_2_^1-122^ was by molecular replacement using a single chain of the HPeV1-2A core domain as a search model (PDB 7ZTW), while LV1-2A_2_ was used as a search model during molecular replacement for SEBV1-2A_2_ in Phaser(25). Phasing for HPeV1-2A^1-129^ by single anomalous dispersion (SAD) made use of the Autosol pipeline in Phenix (26), searching for 3 selenomethionine residues. After density modification, experimental phases were of sufficient quality for automatic model building of 116 of the 129 residues using ARP/wARP (27). This partial model was used as a search model against the native dataset. All models were completed through iterative cycles of model building in COOT (28) and refinement in either Phenix (29), for HPeV3, LV1-2A_2_ and SEBV1-2A_2_, or Refmac5 (30), for LV4-2A_2_ and HPeV1-2A^1-129^, using rigid body, TLS parameter and individual B-factor refinement. At later stages of refinement, we made use of the PDB-REDO server (31) to determine the optimal weights for refinement of all models in Refmac, and the Molprobity server (32) was used to validate the stereochemistry. Data collection and refinement statistics are given in Table 1.

**Table 1.**
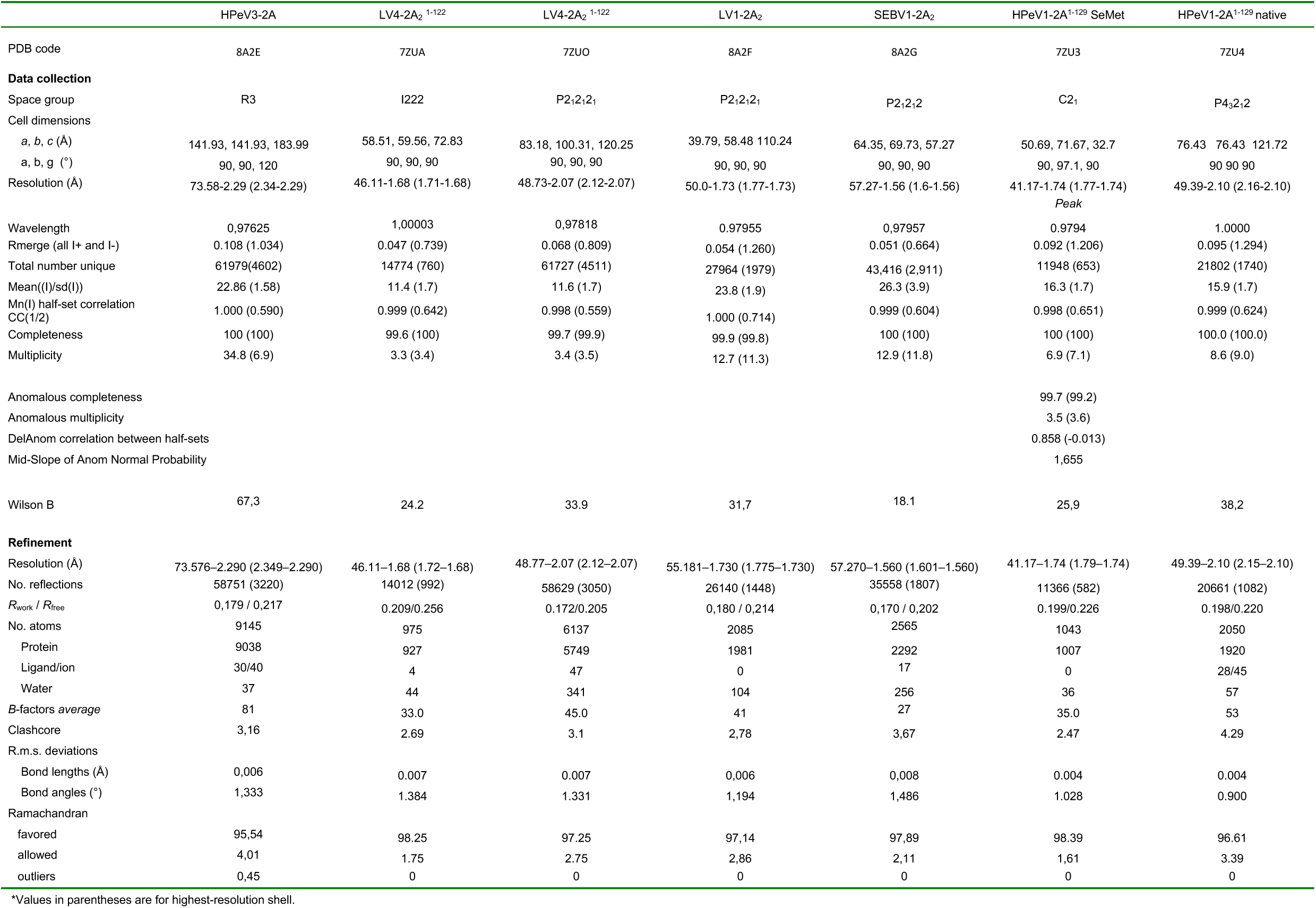
Data collection and refinement statistics.

### 2.4 Small-angle X-ray scattering data collection and processing

SAXS data were recorded at P12 beamline at PETRA-III storage ring (EMBL, DESY, Hamburg, Germany). SEC-SAXS data collection for HPeV1-2A, HPeV3-2A, LV4-2A_2_ and SEBV1-2A_2_ were performed using a Superdex S75 Increase 5/150 column (Cytiva) equilibrated in 20 mM Hepes-NaOH, pH 7.4, 150 mM NaCl, 4 mM TCEP, 1% v/v glycerol, at a flow rate of 0.30-0.35 ml/min (Supplementary Table 1). HPeV1-2A^1-129^ batch samples were dialyzed extensively against buffer containing 20 mM HEPES pH 7.4, 150 mM NaCl, 5 mM DTT and centrifuged for 10 minutes at 12,000 x g before measurement. Measurements were performed in a concentration range of 0.84 – 7.49 mg/ml, experimental details are summarized in Supplementary table 2.

The primary data reduction to obtain 1D scattering profiles was done using the SASFLOW pipeline (33). SEC-SAXS data was further processed using CHROMIXS (34) to produce final background-subtracted SAXS profiles. The analysis of SAXS data was performed using PRIMUS (35) from the ATSAS 2.8 package (36) to obtain the forward scattering, *I*(0), and radius of gyration, *R_g_*, from the Guinier approximation(37) The distribution of pair distances, *p(r)* profile, was calculated with GNOM (38) by Fourier transformation of the data. Concentration-independent molecular weight (MW) estimates were evaluated directly from the SAXS data utilizing Bayesian MW assessment from scattering invariants (39), SAXSMoW (40) and the volume of correlation, Vc(41). All structural parameters and MW estimates are reported in Supplementary Table 1.

The program OLIGOMER (35) was used for fitting of theoretical scattering curves of HPeV1-2A^1-129^ from the monomeric crystallographic structure and derived models of dimer, tetramer and hexamer to the experimental scattering curves measured at eight protein concentrations of in range of 0.84 – 7.49 mg/ml. FFMAKER was applied to create a form-factor file as input and OLIGOMER was run at the maximum scattering vector, *q*_max_, of 4.0 nm^-1^. CRYSOL (42) was used to calculate theoretical scattering profiles from the high-resolution structures and assess the reduced χ² fit to the experimental data. To improve the fit of the high-resolution models to experimental data, crystal structures (as dimers of chains A:B, C:D, E:F and tetramer ABCD for HPeV3-2A) were refined by implementing Normal Mode Analysis (NMA) using program SREFLEX (43). For simplicity we focused on the results for the A:B dimer.

### 2.5 Size-exclusion chromatography coupled with multi-angle laser light scattering (SEC-MALLS)

MALLS experiments were performed following SEC, using a Wyatt Technologies Mini-Dawn TREOS multi-angle light scattering detector coupled to an OptiLab T-Rex refractometer (RI). A Superdex 75 Increase 5/150 analytical column (Cytiva) was equilibrated in 20 mM HEPES-NaOH pH 7.4, 150 mM NaCl, 4 mM TCEP, 1% v/v glycerol, at a flow rate of 0.30-0.35 mL/min, prior to sample injection. The molecular weight (MW) of the protein was determined using the MALLS system at an incident wavelength of 659 nm, combined with the concentration estimates obtained from the RI (dn/dc= 0.185mL/g). The MW distribution from each species eluting from the column was calculated using ASTRA7 software (Wyatt Technology).

## 3. Results and Discussion

### 3.1 Crystal structures of *Parechovirus A-C* H/NC-motif 2A proteins

To determine whether the destroyed active site configuration we observed in the structure of HPeV1-2A (Fig 1A) is unique to this protein or a general feature of *Parechovirus A*, we elucidated the crystal structure of the 2A protein from HPeV3. The structure was determined at 2.29 Å in the *R3* space group with eight molecules in the asymmetric unit and refined to an R_free_ of 21.66%, with 4 Ramachandran outliers (95.54% favored/0.45% outliers), a Molprobity score of 1.71, (97^th^ percentile) and a Ramachandran Z-score of −3.09 ± 0.24 (44) (for crystallographic details see Materials and Methods and Table 1). In all eight chains residues 11-146 could be modeled, and in five of eight chains residue 147 could also be modeled. In chains A, C, E, G crystallographic contacts order the N-terminal residues 1-10, which allowed for their modelling in those chains, while they appear flexible and disordered in the other four chains. In addition, 5 glycerol molecules and 9 sulphate ions as well as 37 waters could be modelled. Overall, the HPeV3-2A monomer adopts a similar structure to HPeV1-2A (r.m.s.d. 4.45 Å for the full chain, 0.85 Å over 97 C_α_ residues of the core identified using Gesamt (45); Fig 1B,F), as expected given their high (88%) sequence identity (Table 2). The N-terminal part of the protein folds into an antiparallel, twisted β-sheet made up of five antiparallel β-strands, where β2 contains the conserved H-box motif. The C-terminal half of the protein, which is predominantly α-helical, exhibits the same rearranged topology as previously observed in HPeV1-2A. The (central) α2-β6-α3 region containing the NC-motif is wrapped around the long α4-helix, which forms the spine of the protein at the back of the twisted β-sheet, thereby positioning the “catalytic” cysteine of the conserved NC-motif on the wrong side of the β-sheet for catalysis. The predicted transmembrane domain (spanning residues 111-129) (18) maps to the α4-helix, and therefore lies at the core of the protein rather than being accessible for membrane insertion/interaction (Fig 1G). At the C-terminus the α5-helix is extended from the globular core of the protein and involved in oligomerization contact formation.

**Figure 1:**
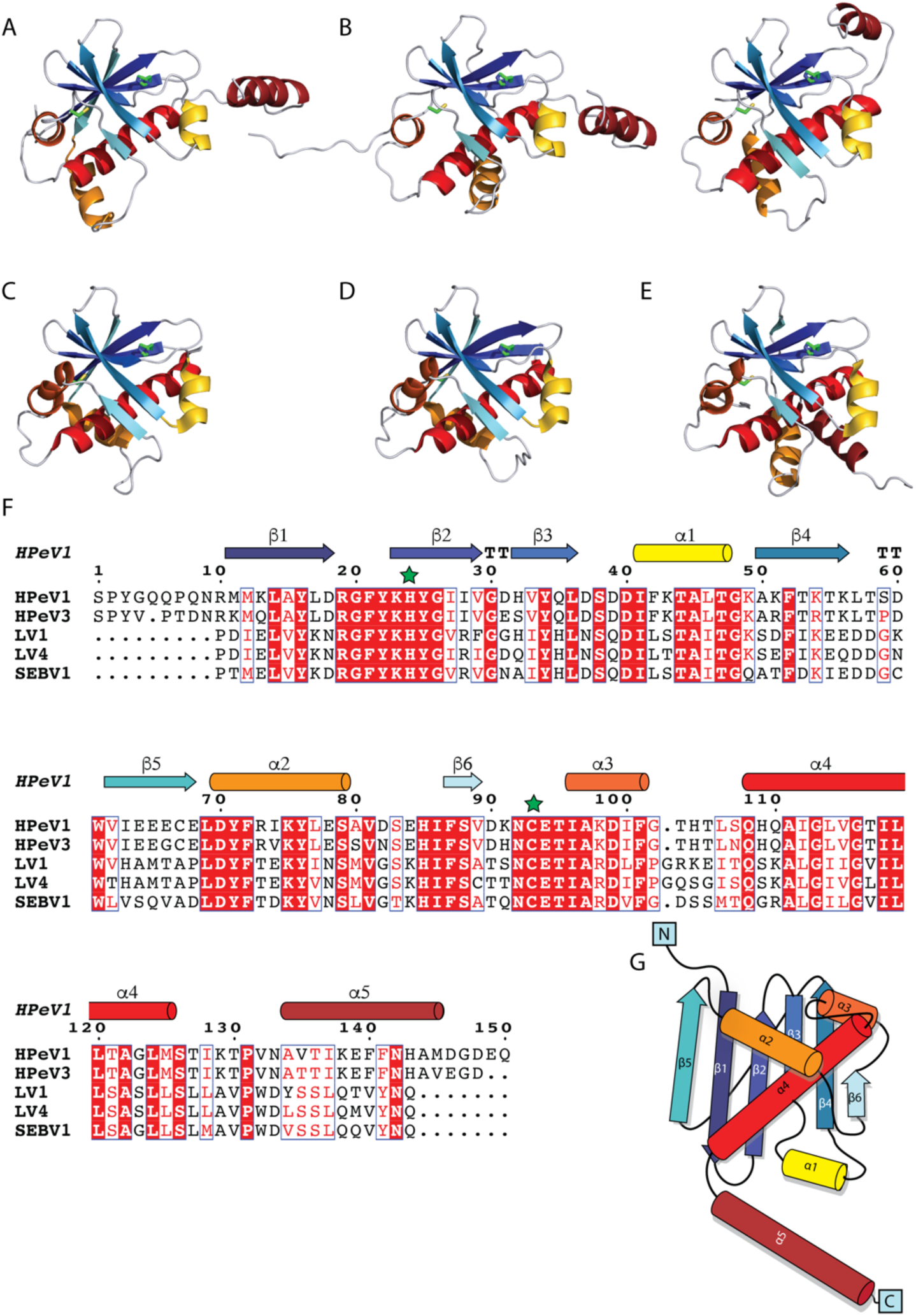
Crystal structures of *Parechovirus* 2A^H/NC^ proteins and their structure-based sequence alignment. All models were aligned to the same orientation and are depicted as cartoon using a modified rainbow coloring scheme with β-strands colored from dark blue to turquoise and α-helices colored from yellow to red from N- to C-terminus. The His and Cys residues of the conserved H-box and NC-motif are depicted as green sticks, highlighting the structural rearrangement destroying the active site configuration. (a) structure of HPeV1-2A, (b) the two distinct conformations of HPeV3-2A observed in the structure, with conformation A on the left and conformation B on the right. (c) structure of LV1-2A2, (d) structure of LV4-2A2^1-122^, (e) structure of SEBV1-2A2, (f) structure-based sequence alignment of *Parechovirus A-C* 2A proteins, generated with ESPript (46), with identical residues shown in white on a red background and similar residues shown in red. The conserved His and Cys are indicated by green stars. Secondary structure elements depicted at the top are taken from the HPeV1-2A structure and colored the same way as in the cartoons of the crystal structures in panels a-e. (g) topology diagram of HPeV1-2A.

**Table 2:** Sequence and structure similarity of *Parechovirus* 2A^H/NC^ proteins. The triangle of the matrix above the identity diagonal shows the sequence identity between the different proteins, while the lower triangle shows r.m.s.d. values between the different structures based on Chimera Matchmaker tool (47), matching the best chain in the reference (the protein in the respective column header) and moving structure for each comparison. The upper number gives the r.m.s.d. and number of paired Cα atoms after pruning, while the lower number gives r.m.s.d and number of Cα atoms for matching of full chains. In the identity diagonal the r.m.s.d between chains of the same protein are listed as calculated by SSM (48)(done separately for conformation A and conformation B in the case of HPeV3).

|  |  | HPeV1 | HPeV3 | LV1 | LV4 | SEBV1 |
| --- | --- | --- | --- | --- | --- | --- |
|  |  | Sequence identity (%) |  |  |  |  |
| HPeV1 | Rmsd (Å) | 0.34 - 0.67 | 86.39 | 44.03 | 46.27 | 46.27 |
| HPeV3* |  | 0.66 (89) | 0.25-0.40 | 42.54 | 44.03 | 45.52 |
|  |  | 4.89 (142) | 0.40-0.69 |  |  |  |
| LV1 |  | 0.77 (92) | 1.00 (81) | 0.72 | 84.44 | 72.39 |
|  |  | 7.29 (128) | 4.34 (125) |  |  |  |
| LV4 |  | 0.91 (95) | 0.98 (82) | 0.66 (115) | 0.64-1.15 | 69.40 |
|  |  | 3.63 (118) | 2.60 (120) | 0.99 (122) |  |  |
| SEBV1 |  | 0.93 (86) | 1.06 (80) | 0.82 (80) | 0.756 (80) | 0.50 |
|  |  | 10.92 (138) | 10.23 (143) | 4.50 (128) | 3.68 (124) |  |

While two monomers of HPeV1-2A assemble to a homodimer and are virtually identical (r.m.s.d. 0.3 Å over 141 C_α_ residues, Fig 2A), the 8 molecules of HPeV3 in the asymmetric unit exhibit some conformational variability and assemble into two homotetramers (Fig 2B). These HPeV3 tetramers are assembled by virtue of the protein adopting two different conformations. The main difference between the two conformations lies in the orientation of the C-terminal helix α5. The first conformation (conformation A) is similar to the conformation of HPeV1 (r.m.s.d. 4.45 Å over the full chain, with the α5 helix rotated by 13° compared to HPeV1; 0.3 Å r.m.s.d. between the four monomers of conformation A), while in conformation B the α4 helix terminates three quarters of a turn later – thereby projecting α5 in a different direction (112-116° rotation outwards away from the helix-exchanging position, with an r.m.s.d. of 7.0 Å over the full chain compared to HPeV1; ∼0.6 Å r.m.s.d. between the four monomers of conformation B). An alternating combination of these two conformations (A-B-A-B pattern) leads to a circular rather than reciprocal exchange of this helix leading to a tetrameric assembly. In addition, the internal region around residues 66-86, corresponding to α2 and its flanking loops, adopts a slightly different orientation, to accommodate this change in oligomerization pattern (discussed in more detail below).

**Figure 2:**
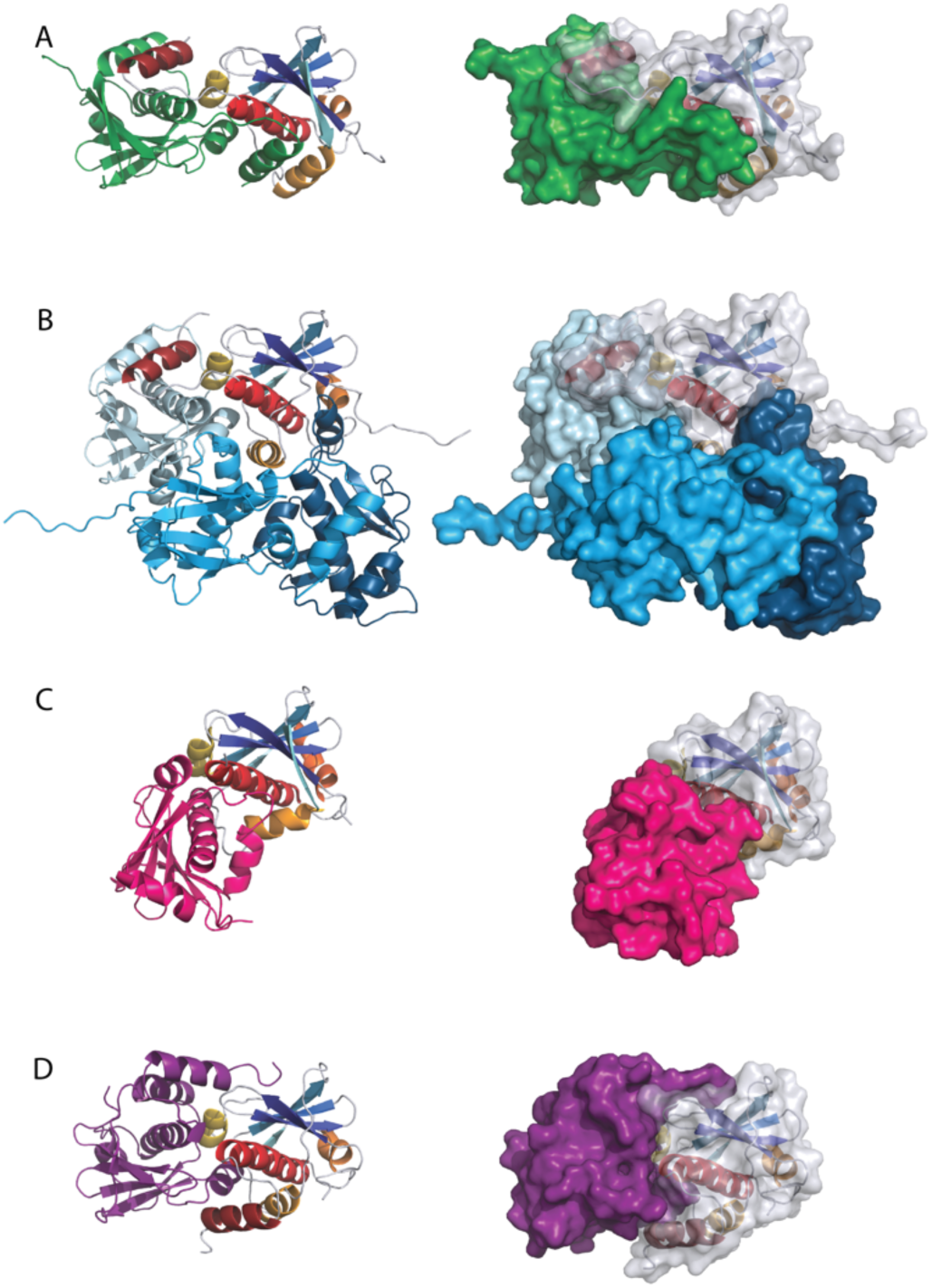
Biological assemblies of *Parechovirus A-C* 2A^H/NC^ proteins. All biological assemblies have been aligned to the same orientation (based on monomer A), and the aligned monomer is depicted in the same cartoon coloring scheme as used in Figure 1. The other monomers in the different biological assemblies have been colored in different colors for the different proteins. The biological assemblies are depicted as cartoon on the left-hand side and as surfaces on the right, with the surface of the aligned monomer semitransparent to allow the cartoon to show through for easier orientation. (a) dimer of HPeV1-2A, (b) tetramer of HPeV3-2A, where the rearrangement of α2 to accommodate the circular tetrameric assembly can be seen in the aligned monomer depicted in modified rainbow coloring (c) dimer of LV4-2A2, (d) dimer of SEBV1-2A2.

To see whether this destroyed active site is also preserved in other species of the genus *Parechovirus*, we decided to expand our structural characterization to members of the *Parechovirus B* species. We determined the structure of LV1-2A_2_ to 1.73 Å resolution in P2_1_2_1_2_1_ with two molecules in the asymmetric unit and refined it to 21.45% Rfree, a Molprobity score of 1.25 (98^th^ percentile) without any Ramachandran outliers and a Ramachandran Z score of 0.06 ±0.61. The final model included 103 water molecules, however, due to missing electron density, the C-terminal residues 130-135 in chain A and residues 123-135 in chain B as well as the C-terminal hexahistidine-tag could not be modeled. We also determined the structure of LV4-2A_2_; interestingly, while the full-length protein did not yield diffraction quality crystals, a C-terminal truncation construct spanning LV4-2A_2_^1-122^ crystallized quite readily and the structure could be determined in two different crystal forms. Crystal form 1 diffracted to 1.68 Å resolution, contained one molecule in the asymmetric unit in space group I222, and was refined to 25.6% R_free_, with no Ramachandran outliers, a Ramachandran Z-score of −1.43±0.89 and a Molprobity score of 1.06 (100^th^ percentile). The final model encompasses residues 1-69 and 74-122, residues 70-73 could not be modelled due to missing electron density. In addition, one acetate ion and 44 water molecules could be modeled. A second crystal form diffracted to 2.1 Å with 6 molecules in the asymmetric unit in space group P2_1_2_1_2_1_ and could be refined to 20.5% R_free_, with no Ramachandran outliers, a Ramachandran Z-score of −1.57± 0.33, a Molprobity score of 1.42 (99^th^ percentile). Due to differences in crystal packing the flexible region spanning residues 69-73 was better ordered, which allowed for their modeling in five of the six chains. Further, the final model also includes 341 water molecules as well as one Tris molecule and several ordered fragments of polyethylene glycol from the crystallization condition (1 EDO, 2 PGE and 2 PEG).

The structures of LV1-2A_2_ (Fig 1C) and LV4-2A_2_ (Fig 1D) are very similar (84% seq.id. and 1.0 Å r.m.s.d. over 121 C_α_ residues). Overall, they exhibit the same topology as HPeV1 (r.m.s.d. 3.13 Å for the full chain; 0.87 Å over the core of 97 C_α_ residues; ∼44% seq. id.; Table 2) and HPeV3 (r.m.s.d. 4.34 Å over the full chain; 0.87 Å over the core of 97 C_α_ residues; ∼44% seq. id; Table 2) (Fig 1F). However, at the N-terminus only the proline of the NPG|P ribosome skipping motif (which separates2A_1_ and 2A_2_) and an aspartate precede β1, compared to 10 residues in HPeV1 (12 residues in HPeV3). Compared to *Parechovirus A* 2A, *Parechovirus B* 2A_2_ also appears to have a slightly shorter C-terminus, which shows an overall greater sequence divergence than the rest of the protein. In our structures this region spanning the potential α5 is either at least partially disordered (LV1) or missing from the construct (LV4) and this leads to some shifts in secondary structure orientation, which also impact its oligomerization. While LV 2A_2_ is also dimeric in the crystal, with the C-terminal α5 helix either missing or at least partially disordered, α2 has shifted to make space and the second molecule is more or less inverted compared to HPeV1-2A, with the second monomer packing onto the face of the protein core where the C-terminal helix (α5) of the second monomer sits in the *Parechovirus A* 2A proteins (discussed in more detail below).

Intrigued by this observed disorder in the C-terminal region of the *Parechovirus B* structures and its apparent influence on protein oligomerization, we decided to extend our structural characterization to the H/NC-motif type 2A_2_ of Sebokelevirus (SEBV1), the sole representative of *Parechovirus C*, which like *Parechovirus B* 2A_2_, exhibits shortened N- and C-termini compared to *Parechovirus A*. The structure of SEBV1-2A_2_ was determined to 1.56 Å resolution in space group P2_1_2_1_2 with two molecules in the asymmetric unit and was refined to 20.16% R_free_, no Ramachandran outliers, a Ramachandran Z-score of −0.40±0.53 and a Molprobity score of 1.16 (98^th^ percentile) (Table 1). All 134 residues could be modeled in both chains, and the final model includes an additional 5 residues at the N- and 7 residues at the C-terminus that were introduced during the cloning procedure, as well one ethylene glycol molecule, a tetraethylene glycol fragment and 256 water molecules. Overall, SEBV1-2A_2_ is similar to the LV 2A_2_ structures (3.70 Å over the full chain; 1.2 Å over the core 97 C_α_ residues), in SEBV1 however, the C-terminal α5 helix is well ordered (and extends to include two of the extra residues past the native protein boundaries) (Fig 1E). Interestingly, while SEBV1-2A_2_ shows the closest sequence identity to LV1-2A_2_ (72%; Table 2), it still shows the same overall dimer organization as *Parechovirus A* 2A (r.m.s.d. 2.21 Å over 224 C_α_ residues within the dimer compared to HPeV1-2A (46% seq. id.); r.m.s.d 2.66 Å over 224 C_α_ residues within the dimer, compared with HPeV3-2A (46% seq. id.)). The r.m.s.d. between full chains is 10.92 Å compared to HPeV1-2A and 10.23 Å for HPeV3-2A; Table 2), except that while the C-terminal α5-helix in HPeV1-2A extends to the second monomer in the dimer, in SEBV1-2A_2_, this helix folds back onto the same monomer and comes to lie in a similar region as the dimer-exchanged helix would occupy. This difference in how α5 is positioned might be due to the fact that in SEBV1-2A_2_ α4 is 4 residues longer, and therefore the following linker towards α5 extends from α4 in a different direction.

### 3.2 Higher order assembly of *Parechovirus* 2A^H/NC^ proteins

Just as the structural rearrangement leading to the destruction of the active site conformation is conserved amongst the *Parechovirus* family members we examined, so is the assembly into dimers, with the exception of HPeV3-2A, which crystallized as homo-tetramers in the asymmetric unit, though these can be seen as a dimer of dimers (Fig 2). It is noteworthy that the interface and configuration of these dimers of the 2A^H/NC^ proteins of HPeV1, LV2 and SEBV1 and the tetramer of HPeV3 all exhibit some differences, and no two proteins exhibit quite the same dimer interface arrangement. While the core domains (excluding α5) of the two molecules in the dimer of HPeV1-2A and SEBV1-2A_2_, and a pair of monomers with conformation A+B in HPeV3-2A adopt the same position and orientation and can be superposed well, the LV-2A_2_ dimer has a shifted arrangement of the two monomers. All three crystal forms of LV1-2A_2_ and LV4-2A_2_ exhibited the same dimeric assembly, confirming that this unusual interface compared to the other family members is not due to crystallographic artefacts, but a characteristic of this species.

We used PISA (49) to analyze the interface contacts and buried surface area of the individual dimers. Across all proteins 12-22% of the accessible surface area participated in interface formation for each monomer and the buried surface area per monomer ranged from 380-1959 Å^2^ (Table S3). Most interface contacts are contributed by the C-terminal α-helical portion of the protein, with a few mapping to the backside of the β-sheet along β5 and β6, if present. Helix α1, which is unique to the *Parechovirus* family members compared to PLAAT3 and AiV-2A is quite centrally positioned in the interface, as is α4, which forms the spine of the protein core at the back of the β-sheet (Fig 1G). The C-terminal α-helix α5, which is another unique feature of the *Parechovirus* 2A^H/NC^ proteins, is also crucial in regulating the oligomeric assembly. In HPeV1 it is exchanged between the two monomers, contributing ∼500 Å^2^ to the buried surface area (Fig 2A). In HPeV3 it is the critical determinant leading to a tetrameric rather than dimeric assembly in the crystal as the two different conformations of α5 in this protein (Fig 1B) lead to a circular rather than reciprocal helix exchange. Monomer A puts α5 onto same place in monomer B as in HPeV1. In monomer B, as a consequence of the extra partial helical turn of α4, there is a ∼112-116° deviation outwards from the dimer axis, projecting α5 onto monomer C (conformation A), capturing the second dimer like a hook. In HPeV3-2A, α2 needs to reorient slightly to accommodate this deviation. In monomer A, α2 is shifted closer to α5 in the helix exchanging dimer seen in HPeV1-2A (to make space for α5 from monomer D), while in monomer B, α2 is in a similar position as in HPeV1-2A, since here α5 from monomer A in the tetramer falls into the same position as in the helix-exchanging dimer arrangement of HPeV1-2A (Fig 2B). The dimer of dimers type assembly is confirmed by PISA (Table S4). While the analysis indicates that higher order assemblies would be stable, the most likely biological assembly is the tetramer seen in the asymmetric unit (ABCD and EFGH), with an average buried surface area (BSA) of 11165 Å^2^ (accessible surface area ∼25500 Å^2^) and a predicted dissociation energy ΔG_diss_≈33.4 kcal/mol. The involved interfaces between monomers A:B and C:D (BSA≈1600 Å^2^), as well as A:D and B:C (BSA≈1010 Å^2^) each have a predicted Complexation Significance Score (CSS) of 1.00 (which would suggest the interface to be stable in solution) and the predicted dissociation pattern is A:B + C:D, in line with the dimers seen for HPeV1-2A.

As mentioned previously, SEBV1-2A_2_ assembles into the same overall dimer configuration as HpeV1, despite having closer sequence identity with LV-2A_2_ (Fig 2D). In SEBV1-2A_2_, the C-terminal helix α5 folds back onto the globular domain, which positions it close to where α2 sits on HPeV1, and α2 shifts ∼7.5Å away from the interface. This folding back of α5 onto the same monomer reduces the dimer interface area somewhat; however, analysis in PISA indicates that this dimer is still stable in solution (CSS=1.00) and the most likely biological assembly, with an interface area of 1595 Å^2^ (compared to ∼1958 Å^2^ for HPeV1-2A), engaging ∼20% of the surface area of each monomer (Tables S3-4). In the SEBV1-2A_2_ dimer an ordered fragment of a PEG molecule lies in a pore formed in the interface, but this is unlikely to have a major influence on dimer formation.

In *Parechovirus B* 2A_2_, the stretch of amino acids corresponding to the C-terminal α5 helix in *Parechovirus A* and *C* 2A proteins, is either at least partially disordered (LV1-2A_2_) or had been truncated to obtain diffraction quality crystals (LV4-2A_2_). Taking a closer look at monomer A of LV1-2A_2_, in which only the last six residues are disordered, we can see that α4 extends for an extra four residues, the same as in SEBV1-2A_2_. However, whereas the remaining C-terminal stretch encompassing α5 in SEBV1-2A_2_ folds back on top of α4 and nestles against α2, in LV1-2A_2_, this stretch falls more to the side of α4 and towards the back of the β-sheet. There, it appears to be anchored by hydrogen bonds formed between main chain carbonyls of the C-terminal stretch with the side chains of Asn9 (β1), Lys14 (β2), and the last ordered residue H-bonds to the side chain hydroxyl of Tyr62 (α2). In conjunction with this different C-terminal organization we also see a different packing of the two monomers forming the dimer, though the majority of the interface contacts are still formed by the α-helices (Fig 2C). SEBV1-2A_2_, HPeV1-2A and HPeV3-2A all assemble into dimers where both monomers have the same relative orientation, with α1 in the middle of the interface, the linker connecting α4 and α5 in each monomer passing each other on the same surface and α5 packing against α2 on opposite faces of the dimer. In LV-2A_2_ on the other hand the second monomer is more or less inverted and helices α2 and α4 occupy the place of α5 in the other *Parechovirus* 2A structures. As a result, both α1 helices abut one another on the edge of the dimer interface, while both α2 helices meet up at the other side of the interface. This dimer configuration leads to the smallest dimer interface with approximately 1000 Å^2^, engaging ∼15% of each monomer’s accessible surface area; however, its CSS score of 1.00 and assembly analysis in PISA indicate that this protein most likely is dimeric, though a tetramer was also found as a possible stable assembly in solution.

To assess whether the oligomeric states observed in the crystal structures (and estimated as stable assembly states by PISA, Table S4) also represented the oligomeric state of the proteins in solution, we performed size-exclusion chromatography (SEC) coupled Multi-Angle Laser Light Scattering (MALLS) and Small Angle X-ray Scattering (SAXS) experiments. In agreement with their dimeric state observed in the crystal, HPeV1-2A, LV4-2A_2_ and SEBV-2A_2_ display a molecular weight of 33.64 kDa, 29.2 kDa and 31.5 kDa, respectively (Fig 3A, C, D, S1), which is within the experimental error of the theoretical molecular weights of the respective dimers (between 30.6 kDa and 34.4 kDa). SEC-SAXS molar mass determination is also in agreement with the values obtained by SEC-MALLS and give a molecular weight range from 25.57-33.82 kDa using the Bayesian inference method (Table S1). However, for HPeV3-2A SEC-MALLS and SEC-SAXS data both reveal that the tetramer observed in the crystal lattice appears to be a crystallization artefact, since SEC-MALLS yields a molecular weight of 38.8 kDa, which is almost twice lower than the theoretical molecular weight of the crystallographic tetramer (71.92 kDa; Fig 3B). SEC-SAXS analysis is also in agreement with a dimeric assembly of HPeV3-2A in solution (28.9 kDa, Table S1). Next, we compared the experimental scattering curve from our SEC-SAXS analyses with the theoretical scattering curve of the crystallographic dimers (A:B, C:D and E:F, or based on crystallographic symmetry when applicable) as well as a tetrameric assembly for HPeV3-2A (ABCD and EFGH), to analyze the size and the shape of each protein in solution (Fig 3, middle panels). The comparison yielded a χ^2^ of 0.99-3.63 for the different dimers and χ^2^ = 72,70 for the ABCD tetramer of HPeV3-2A(Fig 3B, S1), once again inconsistent with tetramerization of this protein in solution. Normal mode analysis, which allows more flexible modeling than Crysol, reduced χ^2^ to 0.94-2.45. These results emphasize once more that these proteins are dimeric in solution.

**Figure 3:**
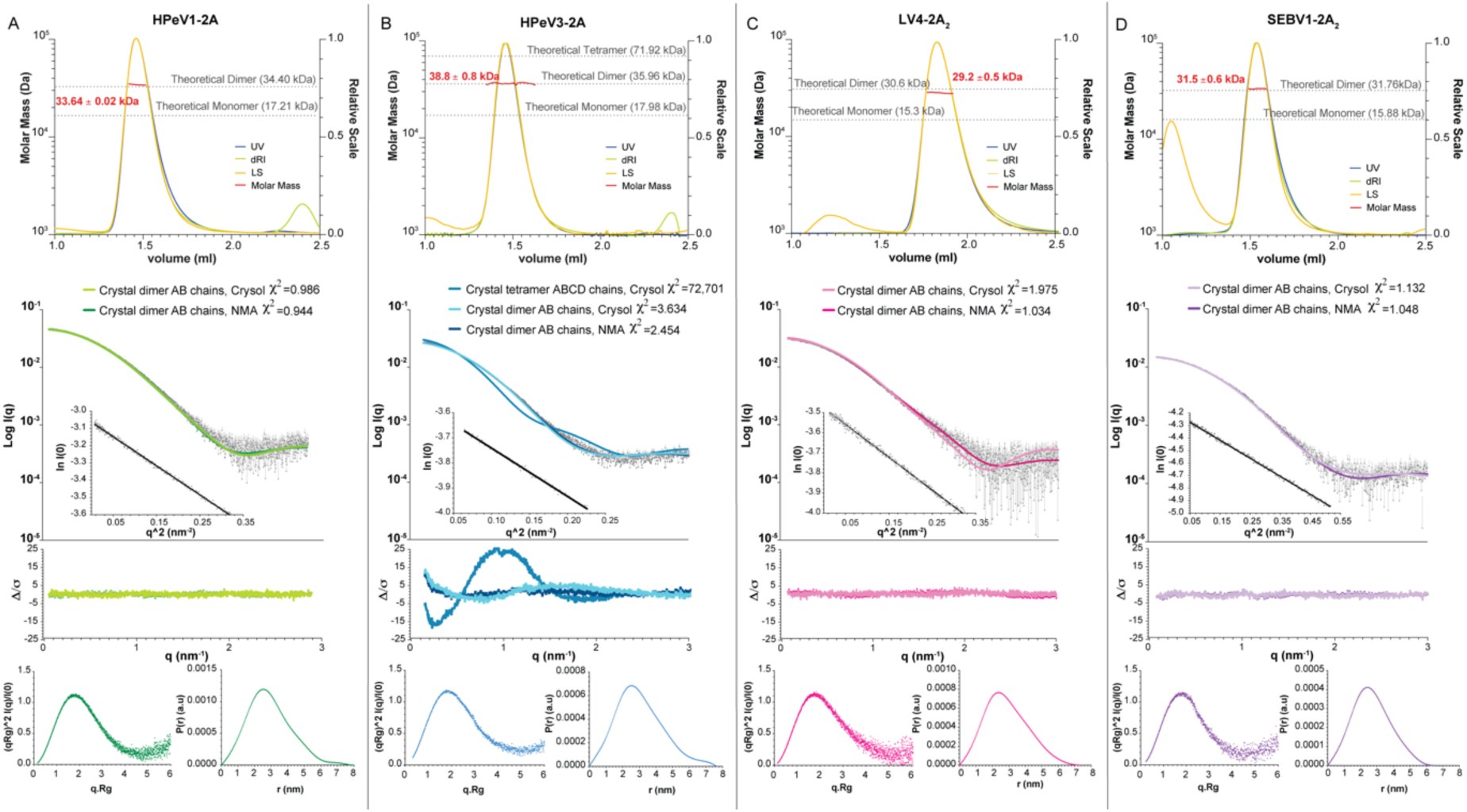
Solution scattering experiments confirm the dimeric assembly of *Parechovirus A-C* 2A^H/NC^ proteins. SEC-MALLS of HPeV1-2A^FL^ (A), HPeV3-2A (B), LV4-2A2 (C) and SeBV1-2A2 (D) allowing the determination of the molecular mass (MW, red) of the protein in solution (top panel). The light scattering (LS, orange), the Abs280nm (blue) and the differential refractive index (dRI, green) are overlaid. The theoretical MW is represented in dashed grey lines. The center of each panel represents experimental SAXS curves of each protein, with the theoretical scattering curves from the high resolutions structures, obtained from rigid body fitting (Crysol) and normal mode analysis (NMA), overlaid on the experimental data. The Guinier region at low angles is represented by the Guinier plot ln I(0) vs q^2^ (inset). On the bottom left of each panel, the dimensionless Kratky plots (qRg)^2^ I(q)/I(0) *vs* q.Rg demonstrate the globular state of the proteins and the interatomic distance distribution P(r) is represented on the bottom right.

The Kratky plots indicate that all the proteins are globular and well-folded. The analysis of the interatomic distance distribution P(r) reveals a D_max_ between 7-8 nm, apart from SEBV-2A_2_ (D_max_ = 6.5 nm). Interestingly, these distances are overall ∼1 nm larger as compared to the respective crystal structures, with the highest difference observed for LV4-2A_2_. Indeed, a ∼ 2 nm difference in D_max_ is observed between the crystal structure and the SEC-SAXS data of LV4-2A_2_. We attribute this discrepancy to the 13 additional residues present in full-length LV4-2A_2_, which had to be truncated to obtain diffraction quality crystals and which presumably exhibit some flexibility around the globular core of the assembly, in agreement with the Kratky plot (Fig 3C) and the missing density for the corresponding residues in the structure of LV1-2A_2_.

### 3.3 Structural plasticity of *Parechovirus* 2A^H/NC^

Given that the C-terminal α5 helix, which appears to be a unique feature in *Parechovirus* 2A^H/NC^ proteins, clearly plays an important role in determining the oligomeric assembly of these 2A proteins, we decided to investigate whether it is indeed the main regulator controlling dimerization.

For LV4-2A_2_ truncation of the C-terminal α5 residues 123-135 does not abrogate oligomerization, as evidenced by the dimeric structure of LV4-2A_2_ in crystals and solution (Fig 2-3, S2).

In the case of HPeV1-2A however, the truncation of the 20 C-terminal residues after α4 had a more profound effect, as HPeV1-2A^1-129^ was no longer an obligate dimer in solution, based on elution volumes during size exclusion chromatography (Fig S2). To see what effect this truncation had on the structure and oligomerization properties of the protein, we determined the crystal structure of the truncated protein, HPeV1-2A^1-129^. A first indication of the profound effect of truncation on the structure came when we were unable to obtain the phases by molecular replacement, despite the availability of a search model with 100% sequence identity and native data to 2.1 Å resolution. Instead, phases were obtained by single-anomalous dispersion method from a selenomethionine-substituted protein crystal diffracting to 1.7 Å resolution in space group C2 with one molecule in the asymmetric unit, followed by automated model building using ARP/WARP ((27)). 116 of the 129 residues could be traced unambiguously in the density of the selenomethionine-substituted protein crystal, and this mostly complete model was used as a molecular replacement model against the native data (space group P4_3_2_1_2 with two molecules in the asymmetric unit). We then completed the model building and refinement of the crystallographic models in both space groups. The final model of the selenomethionine-substituted HPeV1-2A^1-129^ encompassed residues 4-129 and 36 water molecules, had and R_free_ of 22.6%, no Ramachandran outliers, a Ramachandran Z-score of −0.57±0.76 and a Molprobity score of 1.03 (100^th^ percentile). The final model of the native protein was refined to an R_free_ of 22.0%, included residues 8-129 in chain A and residues 12-129 in chain B, as well as nine sulphate ions, four ethylene glycol, two glycerol and 57 water molecules, has no Ramachandran outliers, a Ramachandran Z-score of −0.45±0.58, and a Molprobity score of 1.43 (99^th^ percentile). All three protein chains are similar (r.m.s.d 0.63-0.98 Å between the three different monomers).

To our surprise, the truncated version of HPeV1-2A^1-129^ has undergone significant internal structural rearrangements. While the N-terminal β-sheet is largely unaffected and the C-terminal α4 helix is in the same location packed against the back of the β-sheet, the central region spanning residues 65-107 is rerouted and reverses its directionality, while mostly preserving the secondary structure elements (Fig 4). The conserved NC-motif, which contains the catalytic cysteine in related proteins of the NlpC/P60 superfamily ((50)) is part of the rerouted region, and while the active site configuration is not restored, the rerouting makes the active site Cys more accessible.

**Figure 4:**
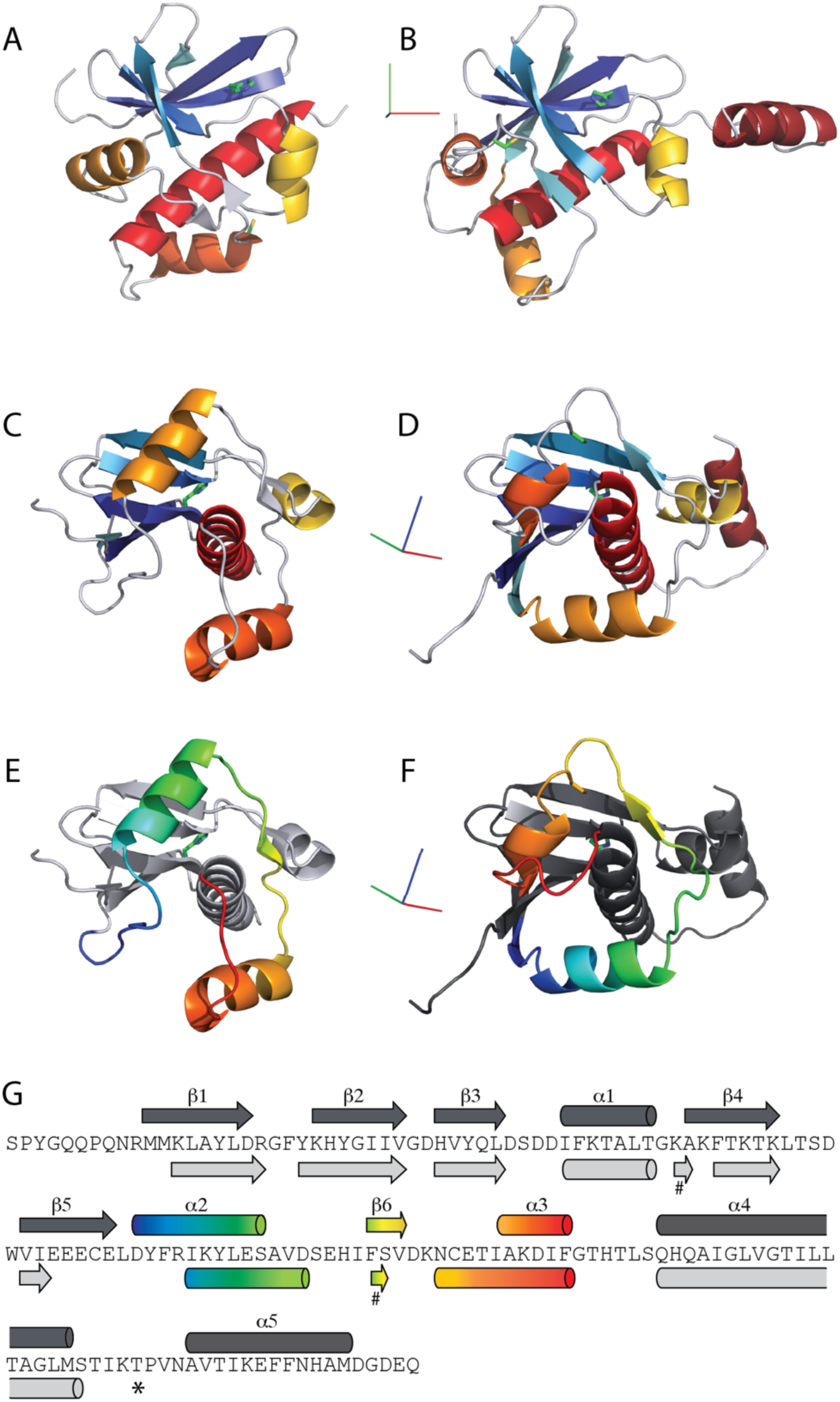
Truncation of the C-terminal oligomerization helix leads to internal topological rearrangements in HPeV1-2A. (a) crystal structure of HPeV1-2A^1-129^, in the same orientation and coloring as used in Figure 1. (b) Cartoon diagram of full-length HPeV1-2A^FL^ in the same orientation, illustrating how the N-terminal β-sheet is preserved as well as the placement of the α4 helix as a spine at the back of the β-sheet. (c-d) HPeV1-2A^1-129^ and HPeV1-2A^FL^ reoriented to visualize the rearranged topology of the α2-β6-α3 region but preserving the modified rainbow coloring of secondary structure elements. (e-f) HPeV1-2A^1-129^ and HPeV1-2A^FL^ in the same orientation as in c-d, but with preserved secondary structure elements colored in grey, and the region with rearranged topology colored in rainbow style from N- to C-terminus. (g) sequence of HPeV1-2A with secondary structure elements of HPeV1-2A^FL^ (above) and HPeV1-2A^1-129^ (below) in the same coloring as in panels e-f. * indicates where HPeV1- 2A^1-129^ has been truncated. In HPeV1-2A^1-129^ β6 participates in a small, secondary β-sheet, indicated by #.

Comparison with the structure of full-length HPeV1-2A^FL^ shows that this rerouting of the central region is not compatible with the dimer conformation observed in HPeV1-2A^FL^, as it induces a steric clash between the rerouted residues 90-92 (adjacent to the NC-motif) and α1 within the unaffected N-terminal half of the protein.

Analysis of crystallographic interfaces for HPeV1-2A^1-129^ reveals a possible alternative dimer in the native protein structure formed by the two molecules in the asymmetric unit with an interface area of ∼1260 Å^2^ and a CSS score of 1.00 (Tables S3-4). In the structure of the selenomethionine substituted protein, there is only one molecule in the asymmetric unit. However, this molecule exhibits an intermolecular disulfide bond in the crystal lattice, engaging its symmetry mate at x+1, y, z+1 (interface area of 1823 Å^2^, CSS = 1.0) (Tables S3-4). This would represent yet another possible dimeric arrangement, distinct from the one seen in either the native HPeV1-2A^1-129^, HPeV1-2A^FL^ or LV-2A_2_ structures. However, it is likely that this latter interface is an artefact induced during crystallization (due to slow oxidation of surface exposed cysteines), and that the different potential dimeric interfaces seen in different crystal forms reveal that this protein no longer is an obligate dimer, due to the absence of α5, a critical determinant for oligomerization behavior. This is also in agreement with the behavior of this truncated protein in solution, where batch-mode SAXS measurements reveal a concentration dependent dynamic monomer-dimer/oligomer equilibrium, exemplified by the linear increase of the observed *I*(0) and *R_g_* (Fig 5A). To quantify the observed concentration-dependent increase in molecular weight, we used OLIGOMER to assess the ratio of oligomeric species in HPeV1-2A^1-129^. Initially we tried to fit the data using only monomers and dimers, however we observed a systematic deviation of the fits at higher concentration. The inclusion of tetramers and hexamers derived from the assembly observed in the crystal structure significantly improved the fits. The analysis showed that at low concentration HPeV1-2A^1-129^ is predominantly monomeric, with the volume fraction close to 90% and small fractions of higher oligomeric assemblies (Fig. 5b). With increasing protein concentration, the fractions of dimers and hexamers become larger. At the highest measured concentration, the OLIGOMER model does not fit the SAXS data, as assessed using CRYSOL (χ² = 1.22, CorMap *P-value* = 0) indicating the presence of other species or structural rearrangement (Fig 5c).

**Figure 5:**
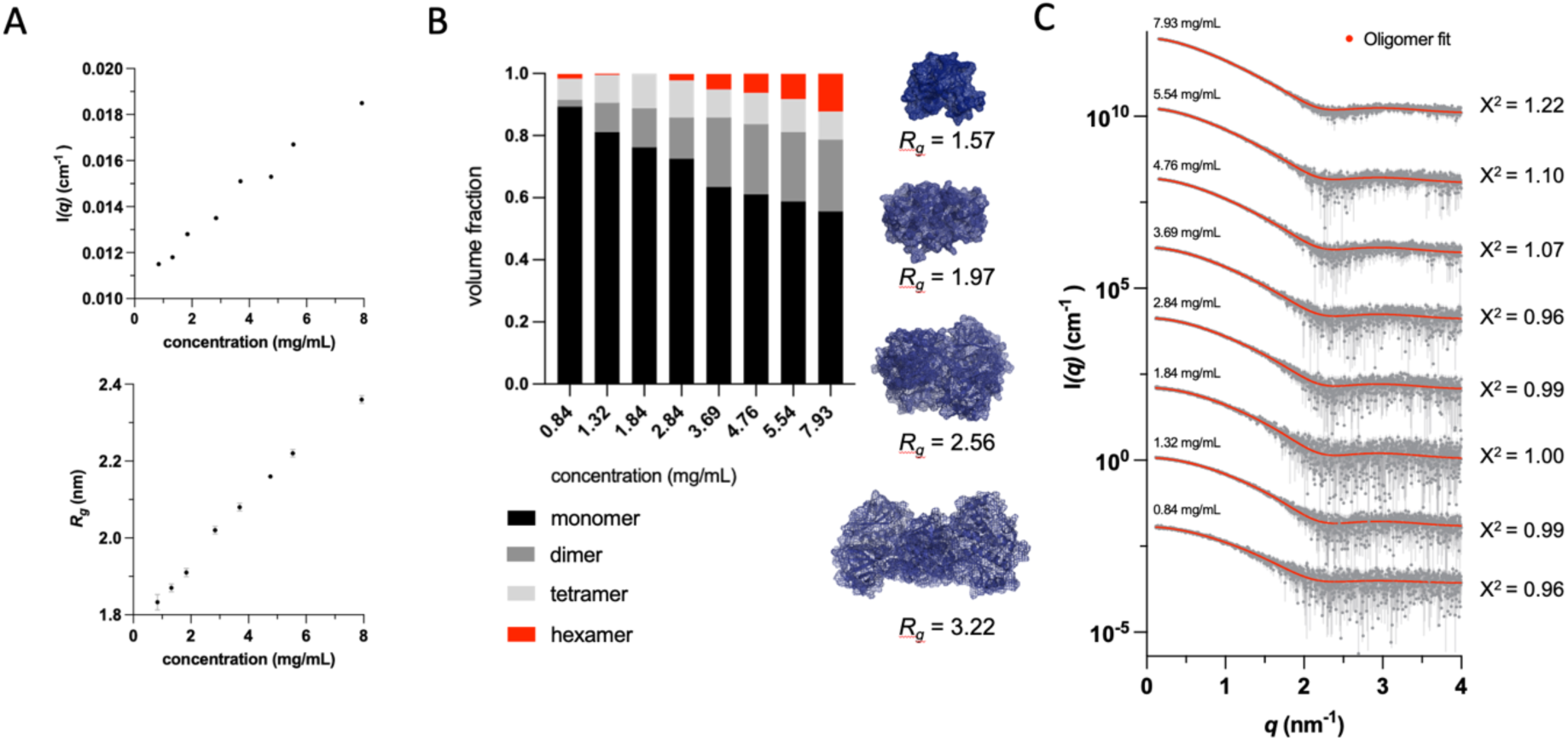
HPeV1-2A^1-129^ is predominantly monomeric in solution, but oligomerizes in a concentration dependent manner. (a) SAXS analysis of HPeV1-2A^1-129^ in batch mode shows *I*(0) (above) and *Rg* (below) increasing with concentration. (b) Stacked column chart visualizing volume fractions of different oligomeric states of HPeV1-2A^1-129^ in solution. Structures of monomer and derived oligomeric species are displayed as mesh and cartoon with theoretical *Rg* values (c) Scattering profiles collected in the concentration range 0.84 - 7.93 mg/mL, experimental data is shown in gray and fits are in red; data shifted along the vertical axis for better visibility.

Analyzing the rerouted topology in the context of the other *Parechovirus* 2A^H/NC^ structures we have elucidated to date, an astonishing structural plasticity is revealed. Comparison of the full-length *Parechovirus* 2A^H/NC^ structures brings to light two main regions of flexibility. The first is the C-terminal α5 helix, which adopts a range of conformations in relation to the globular core of the protein, influencing the oligomerization of the protein. The second apparently flexible stretch centers around α2, the first α-helix after the β-sheet. Its position is “adjustable”, as α2 is basically suspended between anchor points at approximately residues 65 and 87, allowing it to swing out of the way to accommodate different positions of the C-terminal oligomerization helix α5 (Fig. 6). In the truncated HPeV1-2A^1-129^, the flexibility of the α2 region is taken to the extreme by “picking up” the entire internal region between residues 63 within β5 and residue 107 at the beginning of α4, and flipping the directionality of the topology, while preserving the secondary structure elements as well as the position of the protein termini (Fig 4g).

**Figure 6:**
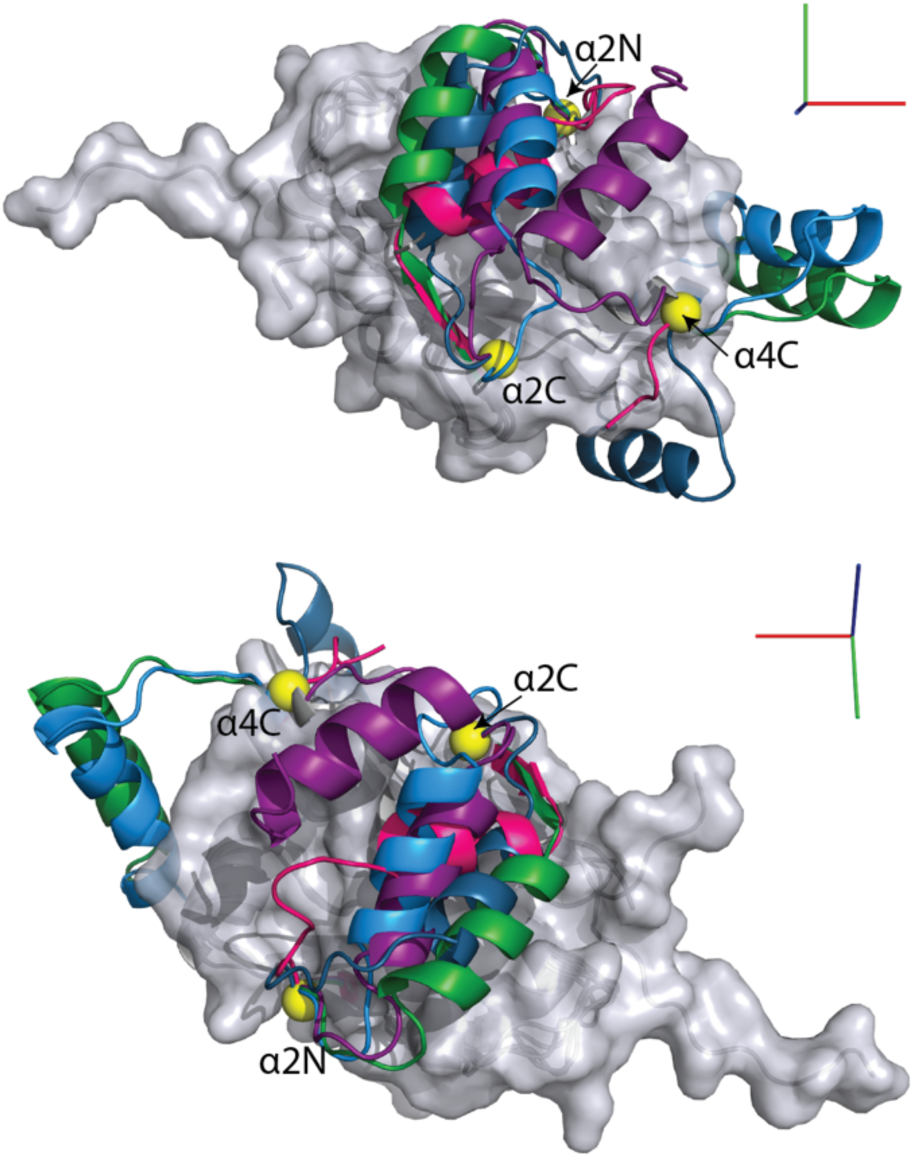
Structural comparison identifies distinct hinge points allowing for the structural plasticity uncovered in *Parechovirus A-C* 2A^H/NC^ proteins. Superposition of the structures of HPeV1-2A, HPeV3-2A (both conformations), LV-2A2 and SEBV1-2A2 as gray cartoon diagrams, with a semitransparent surface displayed for the conserved core of the protein. The flexible secondary structure elements are colored as in Fig. 2: HPeV1-2A in green, HPeV3-2A in blue, LV-2A2 in pink and SEBV1-2A2 in purple. At the end of α4 lies one hinge-point (indicated by a yellow sphere) illustrating the different orientations of the C-terminal oligomerization helix. To accommodate the differences in α5 placement in the different oligomers, the central α2 region, anchored between hinge points α2N and α2C, needs to be mobile and swing out of the way.

## 4. Discussion

Our structural analysis has shown that the topological rearrangement first observed in HPeV1 and leading to the apparent destruction of the active site configuration of the enzyme is indeed a general feature of *Parechovirus* 2A proteins. Furthermore, this study demonstrates that this rearrangement appears to go hand in hand with dimerization of the protein. Could the oligomerization of *Parechovirus* 2A be required for its reported function in RNA binding(17)? Alternatively, could dimerization be required for protein stability after the structural reorganization, since the protein surfaces involved in interface formation in the observed dimers are mostly uncharged or even hydrophobic? One thing that can be excluded is that the protein dimerizes to reconstitute an active site configuration across monomers that is compatible with catalysis.

The C-terminal α5 region, which is a unique C-terminal extension so far only observed in *Parechovirus* 2A proteins, clearly plays a central role in determining the order of oligomerization as well as other correlated and sometimes far-reaching structural rearrangements. Further, the topological rearrangement of HPeV1-2A^1-129^ vs HPeV1-2A^FL^ highlights the importance of correct protein boundaries for the protein in the viral life cycle. As the 2A proteins are produced - and initially fold - as part of the viral polyprotein, the pattern and kinetics of polyprotein processing might influence their structure and oligomeric state in the host cell, which in turn might dictate their function and possible interaction patterns and/or partners. For example, our topologically rearranged HPeV1-2A^1-129^ truncation construct was one of several deletion mutants created by Samuilova and colleagues (17) in their investigation to map which parts of the protein were critical for the RNA interaction they observed in their UV-crosslinking assays. While they could see a strong band corresponding to RNA-cross-linked HPeV1-2A monomers as well as a weaker band corresponding to a cross-linked HPeV1-2A dimer, the truncation to HPeV1-2A^1-129^ resulted in a much weaker band, which was completely lost on truncation to HPeV1-2A^1-107^ or when deleting the internal residues 43-56, encompassing α1. They therefore concluded that the basic-residue rich region 43-56 as well as the C-terminus are important for the RNA-binding protein of the properties. They go on to state that “It seems unlikely that the protein lost its affinity to RNA simply because of an alteration in the conformation of the deletion mutants, but this possibility cannot be completely ruled out.” Our results demonstrate that major conformational changes can occur, and that further experiments are required to pinpoint whether the loss of RNA-interaction observed for HPeV1-2A^1-129^ is because the C-terminal region is required for RNA interaction or because the structural rearrangement disrupts the dimer, which is the functional unit for this protein in the cell.

To fully address how this structural plasticity correlates with the functional repurposing of 2A^H/NC^ in the different picornaviruses we will need an integrative approach combining different structural biology techniques and biophysical methods to explore the conformational dynamics with cell biology and biochemistry to assess the functional implications. This would enable us to recapitulate the possible evolution of this protein from host factor to viral 2A protein with new independent functions.

## Acknowledgements

We thank beamline staff for help during data collection, Cy Jeffries for help with collection and analysis of SAXS data, XXX for critical reading of manuscripts, R. Joosten for help during structure refinement.

## Author Contributions

EvC, LZ, and MP prepared samples; EvC, LZ, MP, ZP, XW and JR performed experiments/assisted in research, EvC, E.F. D.I.S and A.P. designed the study. All authors analysed data. EvC wrote the original draft, EvC, E.F. D.I.S., M.P and A.P. edited and reviewed the manuscript.

## Funding information

This work was supported by funding from the Knut and Alice Wallenberg Foundation (EvC), the Swedish Research Council (EvC), DIS and EEF are funded by UKRI MRC (MR/N00065X/1) and JR by Wellcome (101122/Z/13/Z). LZ was an NDM Prize PhD student. XW is funded by the Institute of Biophysics, Beijing.

---

**Figure S1:**
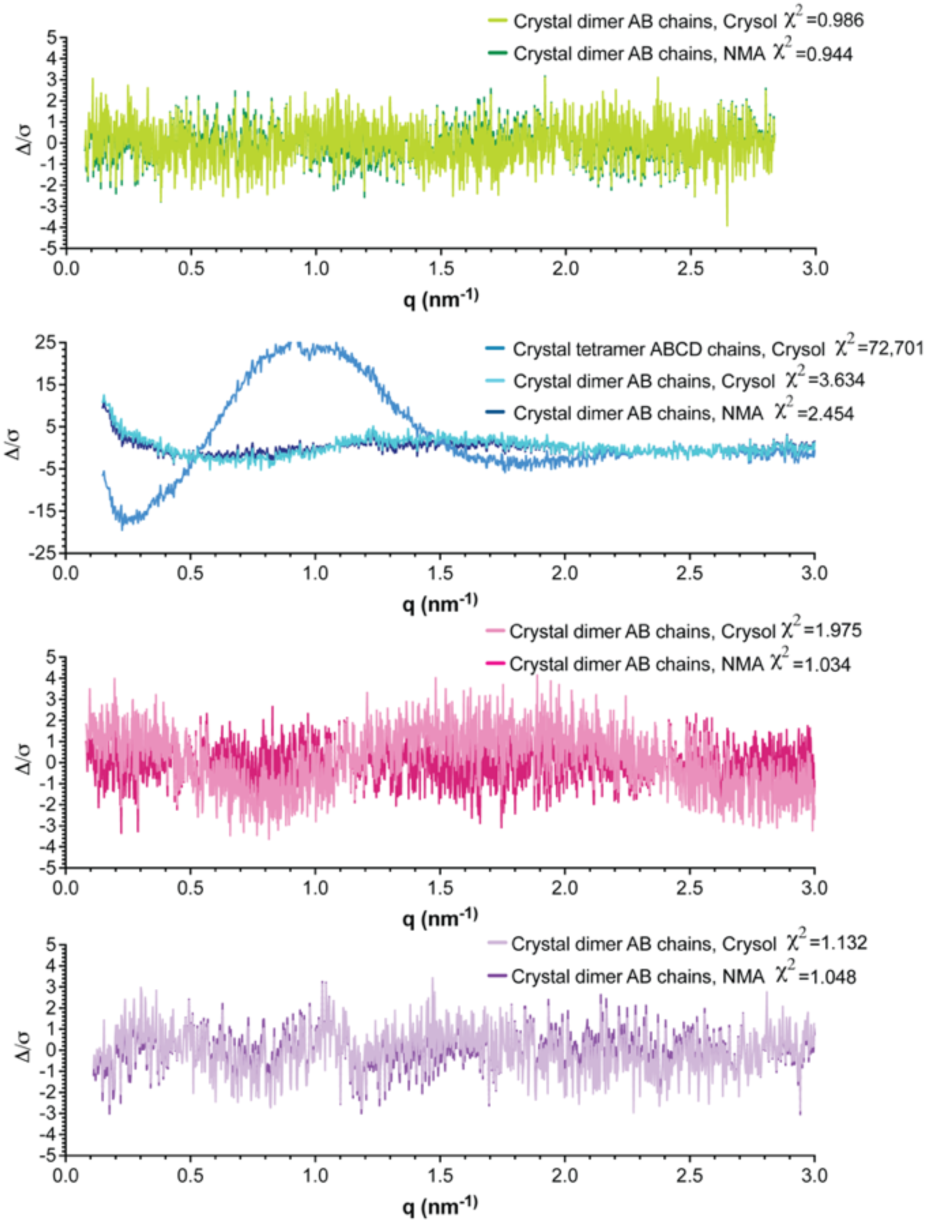
Residuals of Crysol and NMA analyses from Parechovirus A-C SAXS.

**Figure S2:**
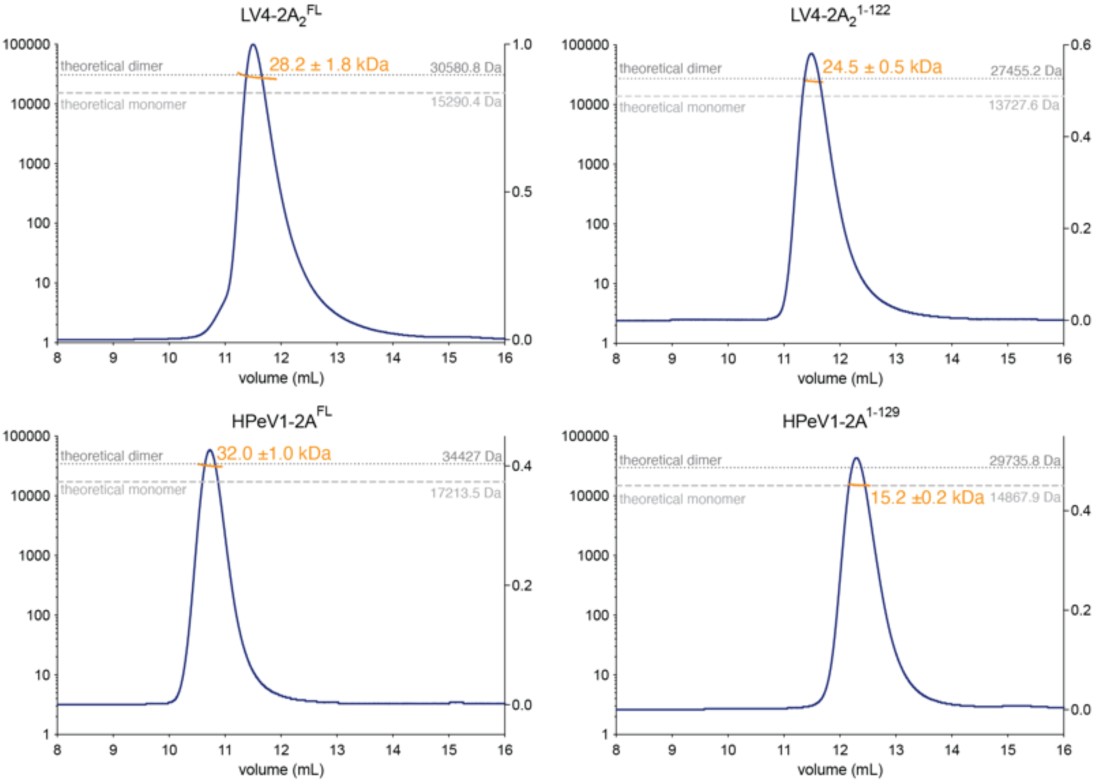
Influence of the truncation of the C-terminal helix on oligomerisation of Parechovirus 2A^H/NC^ proteins. SEC-MALLS analysis of full-length (FL) and truncated LV4-2A2 (A,B) and HPeV1-2A (C,D) with Abs280nm shown in blue with fitted Mw plotted as orange dots across elution peaks. The theoretical Mw of monomeric and dimeric assemblies are plotted as dashed lines.

**Table S1:** SAXS data for *Parechovirus A-C* 2A.

| **Data collection parameters** | **HPeV1-2A FL** | **LV4-2A<sub>2</sub> FL** | **HPeV3-2A** | **SEBV1-2A<sub>2</sub>** |
| --- | --- | --- | --- | --- |
| Instrument | EMBL-P12 (PETRA III, DESY, Hamburg, Germany) | EMBL-P12 (PETRA III, DESY, Hamburg, Germany) | EMBL-P12 (PETRA III, DESY, Hamburg, Germany) | EMBL-P12 (PETRA III, DESY, Hamburg, Germany) |
| Wavelength (nm) | 1.24 | 1.24 | 1.24 | 1.24 |
| Beam size (mm<sup>2</sup>) | 0.2 x 0.05 | 0.2 x 0.05 | 0.2 x 0.05 | 0.2 x 0.05 |
| Detector | Pilatus 6M | Pilatus 6M | Pilatus 6M | Pilatus 6M |
| Sample to detector distance (m) | 3 | 3 | 3 | 3 |
| Exposure time (s)/Number of frames | 1/2880 | 1/2880 | 1/2880 | 1/2880 |
| $q$ range (nm<sup>-1</sup>) | 0.0723 – 2.8361 | 0.08210 – 3.6113 | 0.2264 – 3.3537 | 0.1509 – 3.8735 |
| Temperature (°C) | 25 | 25 | 25 | 25 |
| Type of experiment | SEC-SAXS | SEC-SAXS | SEC-SAXS | SEC-SAXS |
| Column | S75 Increase 5/150 (Cytiva) | S75 Increase 5/150 (Cytiva) | S75 Increase 5/150 (Cytiva) | S75 Increase 5/150 (Cytiva) |
| Injection volume (μl) | 16 | 16 | 30 | 30 |
| Flow rate (ml/min) | 0.35 | 0.3 | 0.35 | 0.35 |
| Concentration (mg/ml) | 10.0 | 10.0 | 10.70 | 8.28 |
| **Structural parameters** |  |  |  |  |
| $I(0)$ (cm<sup>-1</sup>) (from Guinier) | 0.0467 ± 0.00003 | 0.0310 ± 0.00003 | 0.029 ± 0.000042 | 0.0149 ± 0.000023 |
| $R_g$ (nm) (from Guinier) | 2.24 ± 0.07 | 2.21 ± 0.01 | 2.39 ± 0.01 | 2.06 ± 0.01 |
| $sR_g$ range | 0.16 – 1.30 | 0.19 – 1.30 | 0.54 – 1.25 | 0.31 – 1.30 |
| $I(0)$ (cm<sup>-1</sup>) (from $p(r)$) | 0.04674 ± 0.000052 | 0.0310 ± 0.00003802 | 0.02845 ± 0.00003241 | 0.01494 ± 0.00002152 |
| $R_g$ (nm) (from $p(r)$) | 2.262 ± 0.00481 | 2.237 ± 0.003802 | 2.362 ± 0.003289 | 2.068 ± 0.003575 |
| $D_{max}$ (nm) | 8.000 | 7.3000 | 7.63 | 6.47 |
| Porod volume estimate (10<sup>3</sup> nm<sup>3</sup>) | 58.87 | 58.40 | 53824.0 | 31474.0 |
| **Molecular mass determination (kDa)** |  |  |  |  |
| Calculated from amino acid sequence | **17.213** | **15.136** | **17.734** | **15.885** |
| Bayesian inference (probability) | 33.825 (38.14%) | 31.650 (36.74%) | 28.900 (22.22%) | 25.575 (29.94%) |
| Bayesian credibility Interval (probability) | 32.000-34.950 (91.49%) | 28.550-32.000 (97.90%) | 28.550-34.200 (92.19%) | 24.650-28.550 (91.77%) |
| Apparent volume (MoW) | 36.858 | 33.066 | 28.329 | 26.869 |
| Volume of correlation (V<sub>c</sub>) | 34.373 | 30.206 | 32.936 | 27.778 |
| **Software employed for SAXS data reduction and analysis** |  |  |  |  |
| Primary data reduction | SASFLOW | SASFLOW | SASFLOW | SASFLOW |
| Data processing | PRIMUS | PRIMUS | PRIMUS | PRIMUS |
| 3D graphic representation | PyMOL | PyMOL | PyMOL | PyMOL |

**Table S2:** HPeV1-2A^1-129^ SAXS batch measurements.

| *Data collection parameters* |  |
| --- | --- |
| Instrument | EMBL-P12 (PETRA III, DESY, Hamburg, Germany) |
| Wavelength (Å) | 1.24 |
| Beam size (mm<sup>2</sup>) | 0.2 x 0.05 |
| Detector | Pilatus 6M |
| Sample to detector distance (m) | 3 |
| Exposure time (s) | 0.045 |
| $q$ range (Å<sup>-1</sup>) | 0.0097 – 0.48 |
| Temperature (°C) | 10 |
| *Structural parameters* |  |
| Concentration range (mg/mL) | 0.84 – 7.39 |
| I(0) (cm<sup>-1</sup>) (from Guinier) | 0.0115 ± 0.00006 |
| R<sub>g</sub> (nm) (from Guinier) | 1.83 ± 0.017 |
| sR<sub>g</sub> range | 0.21 - 1.30 |
| I(0) (cm<sup>-1</sup>) (from p(r)) | 0.0115 ± 0.00007 |
| R<sub>g</sub> (nm) (from p(r)) | 1.87 ± 0.022 |
| D<sub>max</sub> (nm) | 6.75 |
| Porod volume estimate (nm<sup>3</sup>) | 14.6 |
| Molecular mass determination (kDa) |  |
| Calculated from amino acid sequence | 14.9 |
| I(0) | 15.5 |
| Bayesian inference (probability) | 15.5 (61%) |
| Bayesian credibility Interval (probability) | 14.5-15.8 (91%) |
| Apparent volume (MoW) | 15.7 |
| Volume of correlation (V<sub>c</sub>) | 17.1 |
| **Software employed for SAXS data reduction and analysis** |  |
| Primary data reduction | SASFLOW |
| Data processing | PRIMUS |
| 3D graphic representation | PyMOL |

**Table S3:** PISA interface analysis for the different structures.

| | | Molecule 1 | | | | Molecule 2 | | | | $\Delta iG$ | | $\Delta iG$ | | | | |
| --- | --- | --- | --- | --- | --- | --- | --- | --- | --- | --- | --- | --- | --- | --- | --- | --- |
| PDB |  | Range | iNat | iNres | Surface Å<sup>2</sup> | Range | Symmetry op. | iNat | iNres | Surface Å<sup>2</sup> | Interface area (Å<sup>2</sup>) | kcal/mol | p-value | N hb | N sb | CSS |
| 7ZTW | HPeV1-2A | D | 201 | 51 | 8724 | C | x,y,z | 198 | 50 | 8827 | 1959.2 | -40.9 | 0.004 | 7 | 2 | 1.000 |
|  |  | B | 205 | 50 | 8812 | A | x,y,z | 205 | 52 | 8869 | 1958.2 | -41.2 | 0.003 | 6 | 1 | 1.000 |
|  |  | Average: |  |  |  |  |  |  |  |  | 1958.7 | -41.0 | 0.004 | 7 | 2 | 1.000 |
| 8A2E | HPeV3-2A | D | 163 | 41 | 8762 | C | x,y,z | 169 | 40 | 9724 | 1635.5 | -26.1 | 0.136 | 17 | 2 | 1.000 |
|  |  | H | 161 | 41 | 8746 | G | x,y,z | 165 | 40 | 9552 | 1605.1 | -24.4 | 0.144 | 17 | 2 | 1.000 |
|  |  | B | 159 | 41 | 8664 | A | x,y,z | 167 | 41 | 9676 | 1601.7 | -24.6 | 0.143 | 18 | 3 | 1.000 |
|  |  | F | 158 | 40 | 8675 | E | x,y,z | 166 | 40 | 9609 | 1570.3 | -25.4 | 0.109 | 15 | 3 | 1.000 |
|  |  | Average: |  |  |  |  |  |  |  |  | 1603.2 | -25.1 | 0.133 | 17 | 3 | 1.000 |
|  |  | D | 108 | 25 | 8762 | A | x,y,z | 111 | 32 | 9676 | 1014.0 | -15.3 | 0.194 | 11 | 0 | 1.000 |
|  |  | H | 111 | 26 | 8746 | E | x,y,z | 111 | 32 | 9609 | 1012.3 | -16.5 | 0.162 | 10 | 0 | 1.000 |
|  |  | B | 106 | 23 | 8664 | C | x,y,z | 113 | 30 | 9724 | 1007.3 | -13.5 | 0.318 | 12 | 2 | 1.000 |
|  |  | F | 106 | 24 | 8675 | G | x,y,z | 110 | 32 | 9552 | 992.8 | -14.8 | 0.174 | 11 | 0 | 1.000 |
|  |  | Average: |  |  |  |  |  |  |  |  | 1006.6 | -15.0 | 0.212 | 11 | 1 | 1.000 |
|  |  | C | 48 | 11 | 9724 | A | x,y,z | 46 | 10 | 9676 | 368.0 | -6.3 | 0.300 | 3 | 5 | 0.351 |
|  |  | G | 47 | 11 | 9552 | E | x,y,z | 47 | 10 | 9609 | 360.7 | -4.6 | 0.470 | 2 | 6 | 0.351 |
|  |  | Average: |  |  |  |  |  |  |  |  | 364.3 | -5.5 | 0.385 | 3 | 6 | 0.351 |
| 8A2F | LV1-2A<sub>2</sub> | B | 96 | 25 | 7012 | A | x,y,z | 106 | 30 | 7163 | 1056.9 | -16.1 | 0.107 | 11 | 0 | 1.000 |
| 7ZUO | LV4-2A<sub>2</sub><br>(P212121) | B | 114 | 30 | 6684 | A | x,y,z | 126 | 32 | 6893 | 1204.1 | -27.1 | 0.005 | 7 | 0 | 1.000 |
|  |  | D | 100 | 31 | 6663 | C | x,y,z | 101 | 28 | 6862 | 982.9 | -18.5 | 0.038 | 5 | 0 | 1.000 |
|  |  | F | 77 | 21 | 6896 | E | x,y,z | 66 | 18 | 6329 | 767.8 | -15.7 | 0.027 | 3 | 0 | 1.000 |
|  |  | C | 55 | 17 | 6862 | A | x-1/2,-y-1/2,-z | 47 | 13 | 6893 | 453.7 | -5.0 | 0.451 | 5 | 0 | 0.167 |
|  |  | D | 42 | 13 | 6663 | A | x,y,z | 55 | 18 | 6893 | 450.7 | -3.1 | 0.520 | 5 | 2 | 0.057 |
|  |  | C | 40 | 12 | 6862 | B | x-1/2,y-1/2,-z | 39 | 9 | 6684 | 380.3 | -3.4 | 0.452 | 5 | 1 | 0.133 |
| 7ZUA | LV4-2A<sub>2</sub> | A | 73 | 20 | 6286 | A | x,-y+1,-z+1 | 73 | 20 | 6286 | 725.6 | -11.5 | 0.099 | 6 | 0 | 1.000 |
| 8A2G | SEBV1-2A<sub>2</sub> | B | 163 | 44 | 8068 | A | x,y,z | 169 | 44 | 7877 | 1595.3 | -27.8 | 0.030 | 15 | 0 | 1.000 |
| 7ZU3 | HPeV1-2A<sup>1-129</sup> | A | 160 | 47 | 7929 | A | -x+1,y,-z+1 | 160 | 47 | 7929 | 1823.5 | -46.0 | 0.000 | 9 | 0 | 1.000 |
| 7ZU4 | HPeV1-2A<sup>1-129</sup> | B | 130 | 36 | 7264 | A | x,y,z | 130 | 36 | 7734 | 1259.8 | -28.1 | 0.007 | 6 | 0 | 1.000 |
|  |  | B | 47 | 11 | 7264 | A | -y-1/2,x+1/2,z-1/4 | 48 | 13 | 7734 | 450.1 | -2.7 | 0.517 | 8 | 0 | 0.221 |

**Table S4:** PISA assemblies in the crystal structures of *Parechovirus* 2A protein.

| | Crystal structure | Size | Formula | Composition | Stable | Surface area, (Å<sup>2</sup>) | Buried area, (Å<sup>2</sup>) | $\Delta G_{int}$ (kcal/mol) | $\Delta G_{diss}$ (kcal/mol) |
| --- | --- | --- | --- | --- | --- | --- | --- | --- | --- |
| **7ZTW** | HPeV1-2A | 2 | A<sub>2</sub> | AB | yes | 13760 | 3920 | -41.2 | 34.6 |
|  |  | 2 | A<sub>2</sub> | CD | yes | 13630 | 3920 | -40.9 | 32.0 |
| **8A2E** | HPeV3-2A | 4 | A<sub>2</sub>B<sub>2</sub> | ACBD | yes | 25570 | 11250 | -85.7 | 33.8 |
|  |  | 4 | A<sub>2</sub>B<sub>2</sub> | EGFH | yes | 25500 | 11080 | -85.7 | 33.1 |
|  |  | 2 | AB | CD | yes | 15220 | 3270 | -26.1 | 22.1 |
|  |  | 2 | AB | AB | yes | 15140 | 3200 | -24.6 | 21.2 |
|  |  | 2 | AB | EF | yes | 15140 | 3140 | -25.4 | 20.7 |
|  |  | 2 | AB | GH | yes | 15090 | 3210 | -24.4 | 20.5 |
|  |  | 2 | AB | EH | yes | 16330 | 2020 | -16.5 | 9.4 |
|  |  | 2 | AB | CB | yes | 16370 | 2010 | -13.5 | 7.6 |
|  |  | 2 | AB | AD | yes | 16410 | 2030 | -15.3 | 8.6 |
|  |  | 2 | AB | GF | yes | 16240 | 1990 | -14.8 | 8.1 |
| **7ZUA** | LV4-2A<sub>2</sub> (I222) | 4 | A<sub>4</sub> | A<sub>4</sub> | yes | 20640 | 4500 | -42.4 | 6.8 |
|  |  | 2 | A<sub>2</sub> | A<sub>2</sub> | yes | 11120 | 1450 | -11.5 | 2.9 |
| **8A2F** | LV1-2A<sub>2</sub> | 2 | A<sub>2</sub> | AB | yes | 12060 | 2110 | -16.1 | 9.5 |
| **7ZUO** | LV4-2A<sub>2</sub><br>(P212121) | 6 | A<sub>6</sub> | ABCDEF | yes | 29950 | 10370 | -78.4 | 2.2 |
|  |  | 2 | A<sub>2</sub> | AB | yes | 11170 | 2410 | -27.1 | 18.7 |
|  |  | 2 | A<sub>2</sub> | CD | yes | 11560 | 1970 | -18.5 | 9.3 |
|  |  | 2 | A<sub>2</sub> | EF | yes | 11690 | 1540 | -15.7 | 5.7 |
| **7ZUA** | LV4 2A<sub>2</sub> (I222) | 4 | A<sub>4</sub> | A<sub>4</sub> | yes | 20640 | 4500 | -42.4 | 6.8 |
|  |  | 2 | A<sub>2</sub> | A<sub>2</sub> | yes | 11120 | 1450 | -11.5 | 2.9 |
| **8A2G** | SEBV-1 2A<sub>2</sub> | 2 | A<sub>2</sub> | AB | yes | 12750 | 3190 | -27.8 | 22.4 |
| **7ZU3** | HPV1-2A<sup>1-129</sup> | 2 | A<sub>2</sub> | A<sub>2</sub> | Yes | 12210 | 3650 | -46.0 | 45.9 |
| **7ZU4** | HPeV1-2A<sup>1-129</sup> | 2 | A<sub>2</sub> | AB | yes | 12480 | 2520 | -28.1 | 19.1 |

**Size** – number of monomers participating in assembly; **Formula** – indicates the chemical composition of the assembly. Monomeric units are grouped into chemical types on the basis of their sequence and structure similarity; **Composition** – indicates monomeric units found in the assembly as annotated in the PDB entry; **stable** – contains yes for stable assemblies and no for those which are likely to dissociate in solution. This classification is based on the value of free energy of dissociation; **Surface area** – indicates the solvent-accessible surface area of the assembly (in Å<sup>2</sup>); **Buried area** - indicates the solvent-accessible surface area of monomeric units buried upon assembly formation, (in Å<sup>2</sup>); $\Delta G^{int}$ – indicates the solvation free energy gain upon formation of the assembly, in kcal/M. The value is calculated as difference in total solvation energies of isolated and assembled structures. This value does not include the effect of satisfied hydrogen bonds and salt bridges across the assembly's interfaces; $\Delta G^{diss}$ – indicates the free energy of assembly dissociation, in kcal/M. The free energy of dissociation corresponds to the free energy difference between dissociated and associated states. Positive values of $\Delta G^{diss}$ indicate that an external driving force should be applied in order to dissociate the assembly, therefore assemblies with $\Delta G^{diss}>0$ are thermodynamically stable.

## Notes

### Competing Interest Statement

The authors have declared no competing interest.

## References

1. Zell, R., Delwart, E., Gorbalenya, A. E., Hovi, T., King, A. M. Q., Knowles, N. J., Lindberg, A. M., Pallansch, M. A., Palmenberg, A. C., Reuter, G., Simmonds, P., Skern, T., Stanway, G., Yamashita, T., and Consortium, I. R. (2017) ICTV Virus Taxonomy Profile: Picornaviridae. Journal of General Virology. 98, 2421–2422

2. King, A. M. Q., Brown, F., P., C., T., H., T., H., Knowles, N. J., Lemon, S. M., Minor, P. D., Palmenberg, A. C., Skern, T., and Stanway, G. (1999) Picornaviridae (V., V. R. M. H., M., F. C., L., B. D. H., H., C. C., B., C. E., K., E. M., M., L. S., J., M., A., M. M., J., M. D., R., P. C., and B., and W. R. eds), p. 996, Virus Taxonomy. Seventh Report of the International Committee for the Taxonomy of Viruses., Academic Press, New York, San Diego

3. Coller, B. A., Chapman, N. M., Beck, M. A., Pallansch, M. A., Gauntt, C. J., and Tracy, S. M. (1990) Echovirus 22 is an atypical enterovirus. J Virol. 64, 2692–2701

4. Johansson, S., Niklasson, B., Maizel, J., Gorbalenya, A. E., and Lindberg, A. M. (2002) Molecular analysis of three Ljungan virus isolates reveals a new, close-to-root lineage of the Picornaviridae with a cluster of two unrelated 2A proteins. Journal of Virology. 76, 8920–8930

5. Joffret, M. L., Bouchier, C., Grandadam, M., Zeller, H., Maufrais, C., Bourhy, H., Despres, P., Delpeyroux, F., and Dacheux, L. (2013) Genomic characterization of Sebokele virus 1 (SEBV1) reveals a new candidate species among the genus Parechovirus. Journal of General Virology. 94, 1547–1553

6. Kalynych, S., Pálková, L., and Plevka, P. (2015) The Structure of Human Parechovirus 1 Reveals an Association of the RNA Genome with the Capsid. Journal of Virology. 90, 1377–1386

7. Shakeel, S., Westerhuis, B. M., Domanska, A., Koning, R. I., Matadeen, R., Koster, A. J., Bakker, A. Q., Beaumont, T., Wolthers, K. C., and Butcher, S. J. (2016) Multiple capsid-stabilizing interactions revealed in a high-resolution structure of an emerging picornavirus causing neonatal sepsis. Nature Communications. 7, 11387

8. Domanska, A., Flatt, J. W., Jukonen, J. J. J., Geraets, J. A., and Butcher, S. J. (2019) A 2.8-Angstrom-Resolution Cryo-Electron Microscopy Structure of Human Parechovirus 3 in Complex with Fab from a Neutralizing Antibody. Journal of Virology. 93, 391

9. Zhu, L., Wang, X., Ren, J., Porta, C., Wenham, H., Ekström, J.-O., Panjwani, A., Knowles, N. J., Kotecha, A., Siebert, C. A., Lindberg, A. M., Fry, E. E., Rao, Z., Tuthill, T. J., and Stuart, D. I. (2015) Structure of Ljungan virus provides insight into genome packaging of this picornavirus. Nature Communications. 6, 8316

10. Wang, X., Ren, J., Gao, Q., Hu, Z., Sun, Y., Li, X., Rowlands, D. J., Yin, W., Wang, J., Stuart, D. I., Rao, Z., and Fry, E. E. (2015) Hepatitis A virus and the origins of picornaviruses. Nature Publishing Group. 10.1038/nature13806

11. Lewis-Rogers, N., and Crandall, K. A. (2010) Evolution of Picornaviridae: An examination of phylogenetic relationships and cophylogeny. Molecular Phylogenetics and Evolution. 54, 995–1005

12. 12. Gorbalenya, A. E., and Lauber, C. (2010) Origin and Evolution of the Picornaviridae Proteome. 10.1128/9781555816698.ch16

13. Domanska, A., Guryanov, S., and Butcher, S. J. (2021) A comparative analysis of parechovirus protein structures with other picornaviruses. Open Biol. 11, 210008

14. Danthi, P., Tosteson, M., Li, Q.-H., and Chow, M. (2003) Genome delivery and ion channel properties are altered in VP4 mutants of poliovirus. Journal of Virology. 77, 5266–5274

15. Panjwani, A., Strauss, M., Gold, S., Wenham, H., Jackson, T., Chou, J. J., Rowlands, D. J., Stonehouse, N. J., Hogle, J. M., and Tuthill, T. J. (2014) Capsid Protein VP4 of Human Rhinovirus Induces Membrane Permeability by the Formation of a Size-Selective Multimeric Pore. PLoS Pathog. 10, e1004294

16. Sasaki, J., and Taniguchi, K. (2008) Aichi virus 2A protein is involved in viral RNA replication. Journal of Virology. 82, 9765–9769

17. Samuilova, O., Krogerus, C., Pöyry, T., and Hyypiä, T. (2004) Specific interaction between human parechovirus nonstructural 2A protein and viral RNA. J Biol Chem. 279, 37822–37831

18. Hughes, P. J., and Stanway, G. (2000) The 2A proteins of three diverse picornaviruses are related to each other and to the H-rev107 family of proteins involved in the control of cell proliferation. J. Gen. Virol. 81, 201–207

19. Donnelly, M. L. L., Luke, G., Mehrotra, A., Li, X., Hughes, L. E., Gani, D., and Ryan, M. D. (2001) Analysis of the aphthovirus 2A/2B polyprotein ‘cleavage’ mechanism indicates not a proteolytic reaction, but a novel translational effect: a putative ribosomal ‘skip.’ J Gen Virol. 82, 1013–1025

20. 20. Staring, J., Castelmur, E. von, Blomen, V. A., Hengel, L. G. van den, Brockmann, M., Baggen, J., Thibaut, H. J., Nieuwenhuis, J., Janssen, H., Kuppeveld, F. J. M. van, Perrakis, A., Carette, J. E., and Brummelkamp, T. R. (2017) PLA2G16 represents a switch between entry and clearance of Picornaviridae. Nature. 541, 412–416

21. Luna-Vargas, M. P. A., Christodoulou, E., Alfieri, A., Dijk, W. J. van, Stadnik, M., Hibbert, R. G., Sahtoe, D. D., Clerici, M., Marco, V. D., Littler, D., Celie, P. H. N., Sixma, T. K., and Perrakis, A. (2011) Enabling high-throughput ligation-independent cloning and protein expression for the family of ubiquitin specific proteases. J Struct Biol. 175, 113–119

22. Winter, G. (2009) xia2: an expert system for macromolecular crystallography data reduction. J Appl Crystallogr. 43, 186–190

23. Kabsch, W. (2010) XDS. Acta Crystallogr D Biol Crystallogr. 66, 125–132

24. Evans, P. R., and Murshudov, G. N. (2013) How good are my data and what is the resolution? Acta Crystallogr D Biol Crystallogr. 69, 1204–1214

25. McCoy, A. J., Grosse-Kunstleve, R. W., Adams, P. D., Winn, M., Storoni, L. C., and Read, R. J. (2007) Phaser crystallographic software. J Appl Crystallogr. 40, 658–674

26. Terwilliger, T. C., Adams, P. D., Read, R. J., McCoy, A. J., Moriarty, N. W., Grosse-Kunstleve, R. W., Afonine, P. V., Zwart, P. H., and Hung, L.-W. (2009) Decision-making in structure solution using Bayesian estimates of map quality: the PHENIX AutoSol wizard. Acta Crystallogr Sect D Biological Crystallogr. 65, 582–601

27. Langer, G., Cohen, S. X., Lamzin, V. S., and Perrakis, A. (2008) Automated macromolecular model building for X-ray crystallography using ARP/wARP version 7. Nat Protoc. 3, 1171–1179

28. Emsley, P., and Cowtan, K. (2004) Coot: model-building tools for molecular graphics. Acta Crystallogr D Biol Crystallogr. 60, 2126–2132

29. Liebschner, D., Afonine, P. V., Baker, M. L., Bunkóczi, G., Chen, V. B., Croll, T. I., Hintze, B., Hung, L.-W., Jain, S., McCoy, A. J., Moriarty, N. W., Oeffner, R. D., Poon, B. K., Prisant, M. G., Read, R. J., Richardson, J. S., Richardson, D. C., Sammito, M. D., Sobolev, O. V., Stockwell, D. H., Terwilliger, T. C., Urzhumtsev, A. G., Videau, L. L., Williams, C. J., and Adams, P. D. (2019) Macromolecular structure determination using X-rays, neutrons and electrons: recent developments in Phenix. Acta Crystallogr Sect D Struct Biology. 75, 861–877

30. Murshudov, G. N., Skubák, P., Lebedev, A. A., Pannu, N. S., Steiner, R. A., Nicholls, R. A., Winn, M. D., Long, F., and Vagin, A. A. (2011) REFMAC5 for the refinement of macromolecular crystal structures. Acta Crystallogr D Biol Crystallogr. 67, 355–367

31. Joosten, R. P., Long, F., Murshudov, G. N., and Perrakis, A. (2014) The PDB_REDO server for macromolecular structure model optimization. Iucrj. 1, 213–20

32. Chen, V. B., Arendall, W. B., Headd, J. J., Keedy, D. A., Immormino, R. M., Kapral, G. J., Murray, L. W., Richardson, J. S., and Richardson, D. C. (2010) MolProbity: all-atom structure validation for macromolecular crystallography. Acta Crystallogr D Biol Crystallogr. 66, 12–21

33. Franke, D., Kikhney, A. G., and Svergun, D. I. (2012) Automated acquisition and analysis of small angle X-ray scattering data. Nucl Instruments Methods Phys Res Sect Accel Spectrometers Detect Assoc Equip. 689, 52–59

34. Panjkovich, A., and Svergun, D. I. (2018) CHROMIXS: automatic and interactive analysis of chromatography-coupled small-angle X-ray scattering data. Bioinformatics. 34, 1944–1946

35. Konarev, P. V., Volkov, V. V., Sokolova, A. V., Koch, M. H. J., and Svergun, D. I. (2003) PRIMUS: a Windows PC-based system for small-angle scattering data analysis. J Appl Crystallogr. 36, 1277–1282

36. Franke, D., Petoukhov, M. V., Konarev, P. V., Panjkovich, A., Tuukkanen, A., Mertens, H. D. T., Kikhney, A. G., Hajizadeh, N. R., Franklin, J. M., Jeffries, C. M., and Svergun, D. I. (2017) ATSAS 2.8: a comprehensive data analysis suite for small-angle scattering from macromolecular solutions. J Appl Crystallogr. 50, 1212–1225

37. Guinier, A. (1939) La diffraction des rayons X aux très petits angles: application à l’étude de ph́enom̀enes ultramicroscopiques. Ann. Phys. 11, 161–237

38. Svergun, D. I. (1992) Determination of the regularization parameter in indirect-transform methods using perceptual criteria. J Appl Crystallogr. 25, 495–503

39. Hajizadeh, N. R., Franke, D., Jeffries, C. M., and Svergun, D. I. (2018) Consensus Bayesian assessment of protein molecular mass from solution X-ray scattering data. Sci Rep-uk. 8, 7204

40. 40. Fischer, H., Neto, M. de O., Napolitano, H. B., Polikarpov, I., and Craievich, A. F. (2010) Determination of the molecular weight of proteins in solution from a single small-angle X-ray scattering measurement on a relative scale. J Appl Crystallogr. 43, 101–109

41. Rambo, R. P., and Tainer, J. A. (2013) Accurate assessment of mass, models and resolution by small-angle scattering. Nature. 496, 477–481

42. Svergun, D., Barberato, C., and Koch, M. H. J. (1995) CRYSOL – a Program to Evaluate X-ray Solution Scattering of Biological Macromolecules from Atomic Coordinates. J Appl Crystallogr. 28, 768–773

43. Panjkovich, A., and Svergun, D. I. (2015) Deciphering conformational transitions of proteins by small angle X-ray scattering and normal mode analysis. Phys Chem Chem Phys. 18, 5707–5719

44. Sobolev, O. V., Afonine, P. V., Moriarty, N. W., Hekkelman, M. L., Joosten, R. P., Perrakis, A., and Adams, P. D. (2020) A Global Ramachandran Score Identifies Protein Structures with Unlikely Stereochemistry. Struct Lond Engl 1993. 28, 1249–1258.e2

45. Krissinel, E. (2012) Enhanced fold recognition using efficient short fragment clustering. J Mol Biochem. 1, 76–85

46. Robert, X., and Gouet, P. (2014) Deciphering key features in protein structures with the new ENDscript server. Nucleic Acids Res. 42, W320–W324

47. Meng, E. C., Pettersen, E. F., Couch, G. S., Huang, C. C., and Ferrin, T. E. (2006) Tools for integrated sequence-structure analysis with UCSF Chimera. Bmc Bioinformatics. 7, 339–339

48. Krissinel, E., and Henrick, K. (2004) Secondary-structure matching (SSM), a new tool for fast protein structure alignment in three dimensions. Acta Crystallogr Sect D Biological Crystallogr. 60, 2256–2268

49. Krissinel, E., and Henrick, K. (2007) Inference of macromolecular assemblies from crystalline state. J Mol Biol. 372, 774–797

50. Anantharaman, V., and Aravind, L. (2003) Evolutionary history, structural features and biochemical diversity of the NlpC/P60 superfamily of enzymes. Genome Biol. 4, R11

